# Small-molecule inhibitor of C-terminal HSP90 dimerization modulates autophagy and functions synergistically with mTOR inhibition to kill cisplatin-resistant cancer cells

**DOI:** 10.1101/2024.12.16.628644

**Authors:** Céline David, Yadong Sun, Vitalij Woloschin, Melina Vogt, Niklas Dienstbier, David Schlütermann, Lena Berning, Beate Lungerich, Annabelle Friedrich, Seda Akgün, María José Mendiburo, Christopn G.W. Gertzen, Arndt Borkhardt, Sebastian Wesselborg, Holger Gohlke, Sanil Bhatia, Thomas Kurz, Björn Stork

## Abstract

**Background:** A major obstacle for the successful treatment of cancer is the primary presence or development of resistance mechanisms toward therapeutic intervention. In urothelial cancer, cisplatin-based regimens are still routinely employed, and multiple cellular pathways contribute to chemoresistance. Since the identification of heat shock protein 90 (HSP90) as potential cancer target, various HSP90 inhibitors (HSP90i) have been developed and evaluated in clinical trials. However, limited efficacy has been observed, mainly caused by dose-limiting toxicity and the concomitant induction of a cytoprotective heat shock response (HSR). To avoid this effect, inhibitors targeting the C-terminal domain (CTD) of HSP90 that do not elicit an HSR have been put forward. Additionally, the crosstalk between autophagy and HSP90 is currently being explored, since both processes work together in proteostasis, and the modulation of autophagic responses might be helpful in order to improve the efficacy of HSP90 inhibitors.

**Methods:** The second-generation small-molecule inhibitor VWK147 targeting HSP90 CTD dimerization was synthesized and characterized in detail by biochemical cell-free and cellular assays and molecular modeling. Specifically, HSP90 inhibition, cell viability, and autophagy were monitored in mono- and combined treatments.

**Results:** We demonstrate that VWK147 induces cell death in both cisplatin-sensitive and cisplatin-resistant urothelial carcinoma cells. The treatment with VWK147 in these cells led to the destabilization of classical HSP90 client proteins without triggering an HSR. Additionally, we observe that VWK147 re-sensitizes resistant urothelial carcinoma cells to cisplatin and—in combination with mTOR inhibition—synergistically kills cisplatin-sensitive and -resistant cells, in contrast to what is observed upon treatment with the N-terminal domain-targeting HSP90 inhibitor 17-AAG. This synergy may be explained by VWK147-mediated inhibition of late autophagy events, and thus a blockade of autophagic flux. Finally, we also observed that VWK147 induces non-canonical LC3 lipidation, indicating that this compound possibly exerts a broader effect on ion balance or pH of the endolysosomal system.

**Conclusion:** VWK147 is a promising inhibitor that targets the C-terminal dimerization of HSP90 and simultaneously exhibits autophagy-modulating effects. This compound could potentially be an effective option for improving anti-cancer therapies and/or overcoming treatment resistance.

## Background

Heat shock proteins (HSPs) represent a large family of molecular chaperones ensuring the function and stability of proteins. The major groups of HSPs are subgrouped based on their molecular weight (1). The HSP90 family includes five members, with either cytoplasmic (HSPC1/HSP90AA1; HSPC2/HSP90AA2; HSPC3/HSP90AB1), endoplasmic reticulum (HSPC4/HSP90B1), or mitochondrial localization (HSPC5/TRAP1) (2). The general structure of HSP90 proteins includes 1) an N-terminal domain (NTD) harboring the ATPase activity and mediating co-chaperone binding, 2) a middle domain (MD) mediating client protein binding, co-chaperone binding and ATPase activation, and 3) a C-terminal domain (CTD) required for dimerization and co-chaperone binding (1). The expression of heat shock proteins is upregulated in response to several cellular stress conditions, including raised temperature or oxidative stress. Under these conditions, HSPs support the recovery or elimination of denatured or aggregated proteins (3). HSP90 promotes cancer development by regulating several hallmarks of cancer, including sustained proliferative signaling, evasion of growth inhibitors, activation of invasion and metastasis, induction of angiogenesis, or resistance to cell death (1, 4). Furthermore, HSP90 is overexpressed in several cancers, and high HSP90 expression has been identified as a marker of poor prognosis in lung cancer, esophageal cancer, urinary bladder cancer, melanoma, and leukemia (1). Accordingly, HSP90 became an attractive target for anti-cancer therapy already in the early 1990’s (5). So far, over 19 inhibitors targeting the HSP90 NTD have entered clinical trials, but none have been approved by the FDA as anti-cancer monotherapy (5). Recently, the HSP90 NTD targeting inhibitor TAS-116 (Pimitespib) has been approved by the Japanese FDA for the treatment of gastrointestinal stromal tumors refractory to standard tyrosine kinase inhibitors (6). Nevertheless, major disadvantages of NTD-targeting inhibitors are dose-limiting toxicities and the induction of a heat shock response (HSR) (5), during which HSP70 and other HSR indicators become upregulated, ultimately leading to cytoprotection of cancer cells and to increased resistance to HSP90 inhibition-induced cell death (7). Next to NTD inhibitors, isoform-selective and CTD inhibitors have been described (5). CTD-targeting inhibitors bind to the C-terminal ATP-binding site (e.g., novobiocin and derivatives), to co-chaperone binding sites, or interfere with HSP90 dimerization (5, 8–10). With regard to dimerization-inhibiting compounds, peptidic inhibitors, a peptidomimetic inhibitor (aminoxyrone), and a small molecule inhibitor (5b or LSK82) based on a tripyrimidonamide scaffold have previously been reported (8–10). In this study, we report the synthesis and evaluation of the second-generation small-molecule CTD dimerization inhibitor VWK147.

Generally, HSP90 exerts its chaperone function by forming complexes with the co-chaperone and the client in order to ensure their maturation and/or to maintain their stability (11, 12). In turn, cellular proteostasis is mainly regulated by the ubiquitination-proteasome system (UPS) and (macro-)autophagy pathways, and both pathways contribute to the degradation of HSP90 clients (11). However, autophagy not only mediates the degradation of HSP90 clients, but HSP90 associates and regulates several of the autophagy effector proteins. During the process of autophagy, proteins and organelles are transferred to lysosomes, where they become degraded and recycled. The initiation of autophagy is regulated by the ULK1 protein kinase complex and the VPS34/PIK3C3 lipid kinase complex. The core ULK1 complex consists of the Ser/Thr kinase unc-51 like autophagy activating kinase 1 (ULK1) and the associated proteins autophagy-related 13 (ATG13), ATG101, and focal adhesion kinase family interacting protein of 200 kD (FIP200) (13). The ULK1 complex is regulated by the mammalian target of rapamycin (mTOR), which suppresses ULK1 kinase activity and thus inhibits autophagy via the phosphorylation of ULK1 at Ser758 (equivalent to Ser757 of mouse ULK1) (14, 15). This site becomes dephosphorylated upon starvation or the pharmacological inhibition of mTOR, ultimately leading to ULK1 activation and the induction of autophagy (14, 15). The second autophagy-inducing complex, the VPS34/PIK3C3 complex, is a class III phosphoinositide 3-kinase (PI3K) and contains the catalytic subunit vacuolar protein sorting 34 (VPS34)/phosphatidylinositol 3-kinase catalytic subunit type 3 (PIK3C3) and the associated proteins phosphoinositide-3-kinase regulatory subunit 4 (PIK3R4), ATG14, Beclin 1, and nuclear receptor binding factor 2 (NRBF2) (16). The product of the VPS34/PIK3C3-catalyzed reaction, phosphatidylinositol 3-phosphate (PI3P), then recruits further downstream effector proteins such as double FYVE-containing protein 1 (DFCP1) and WD-repeat protein interacting with phosphoinositides (WIPI) proteins (17). WIPI2 mediates the recruitment of the ATG12— ATG5-ATG16L1 complex, which is required for the lipidation of ATG8 proteins such as [microtubule associated protein 1] light chain 3 ([MAP1]]LC3) (17). During selective autophagy, ATG8 proteins bind to LC3-interacting region (LIR) motifs of the cargo or autophagy-receptors such as sequestosome 1 (SQSTM1)/p62 (18). Next to canonical autophagy, non-canonical pathways have also been identified, which are characterized by the conjugation of ATG8s to single membranes (CASM) (19, 20). On the molecular level, CASM appears to depend on the increased association of the lysosomal V-ATPase V0–V1 subunits, facilitating ATG16L1 recruitment through its C-terminal WD40 domain (21). Thus, inhibition of the lysosomal V-ATPase with bafilomycin A_1_ (BafA_1_) leads to increased levels of lipidated LC3 during canonical autophagy, but inhibits LC3 lipidation during CASM (19).

In the last two decades, the function of autophagy in cancer has been addressed by various studies. It appears that autophagy plays an ambivalent role in cancer, i.e. prevention of early tumorigenesis versus the support of survival of established and metastasizing tumors (22). Additionally, autophagic processes in the tumor microenvironment and/or immune-relevant cells, and non-canonical autophagy pathways such as CASM have to be considered in order to understand the role of autophagy in cancer pathogenesis (22). Nevertheless, the identification and characterization of specific autophagy-modulating small molecules remains a promising approach for cancer therapy.

Urinary bladder cancer (BC) is the 9th most common cancer worldwide, with 613,791 new cases and 220,349 deaths in 2022 (23). With 90% of these cases, urothelial carcinoma (UC) is the most common histologic type of BC (24). Cisplatin-based combination treatments still represent an important adjuvant therapy, but their application and efficacy are frequently limited by the development of chemoresistance (25). In bladder cancer, the PI3K/AKT/mTOR pathway is constitutively activated in up to 40% of tumors (26). Furthermore, Pan et al. report an overexpression of both HSP90 and mTOR in bladder cancer cells (27). Here, we demonstrate that the small molecule VWK147 induces cell death in cisplatin-sensitive and -resistant UC cells (UCCs), and re-sensitizes resistant cells to cisplatin treatment. In these UCCs, VWK147 treatment leads to destabilized client proteins without a concomitant heat shock response. Furthermore, we observe that VWK147—but not the N-terminal domain-targeting HSP90 inhibitor 17-AAG—in combination with the mTOR inhibitor Torin2 synergistically kills cisplatin-sensitive and -resistant UCCs. Finally, we show that VWK147 induces non-canonical LC3 lipidation but inhibits late stages of canonical autophagy.

### Results

### Synthesis of VWK147 as CTD HSP90 inhibitor

Drawbacks of the previously described α-helix mimetic **1** (5b or LSK82, Figure 1A, (9)) are the only moderate anti-cancer activity and its quite poor water solubility (aq. solubility). To simplify the chemical structure of **1** (LSK82) and to find out whether the benzoyl cap group is required for HSP90 inhibition and anti-cancer activity, we aimed at removing the benzoyl cap group of **1** to provide **2** (VWK147, Figure 1A). Tripyrimidonamide **1** (LSK82) was synthesized as starting material from monomeric building blocks using a modular approach according to our previously published protocols (9, 28). For the preparation of **2** (VWK147), the benzoyl cap group of **1** was removed by heating of **1** in a diluted solution of sodium hydroxide in methanol (Figure 1). The side chains of the resulting cap-less tripyrimidonamide **2** (VWK147) mimic the same hot spots (Y689, I692, L696) of the CTD dimerization interface of HSP90 as **1**. The dimerization hot spots are located on α-helix H5, form a functional epitope, and account for most of the HSP90 dimerization energy. Since the C-terminal dimerization is essential for the chaperone activity of HSP90, in the following, **2** (VWK147) was studied by molecular modeling as well as in biochemical cell-free and cellular assays.

**Figure 1:**
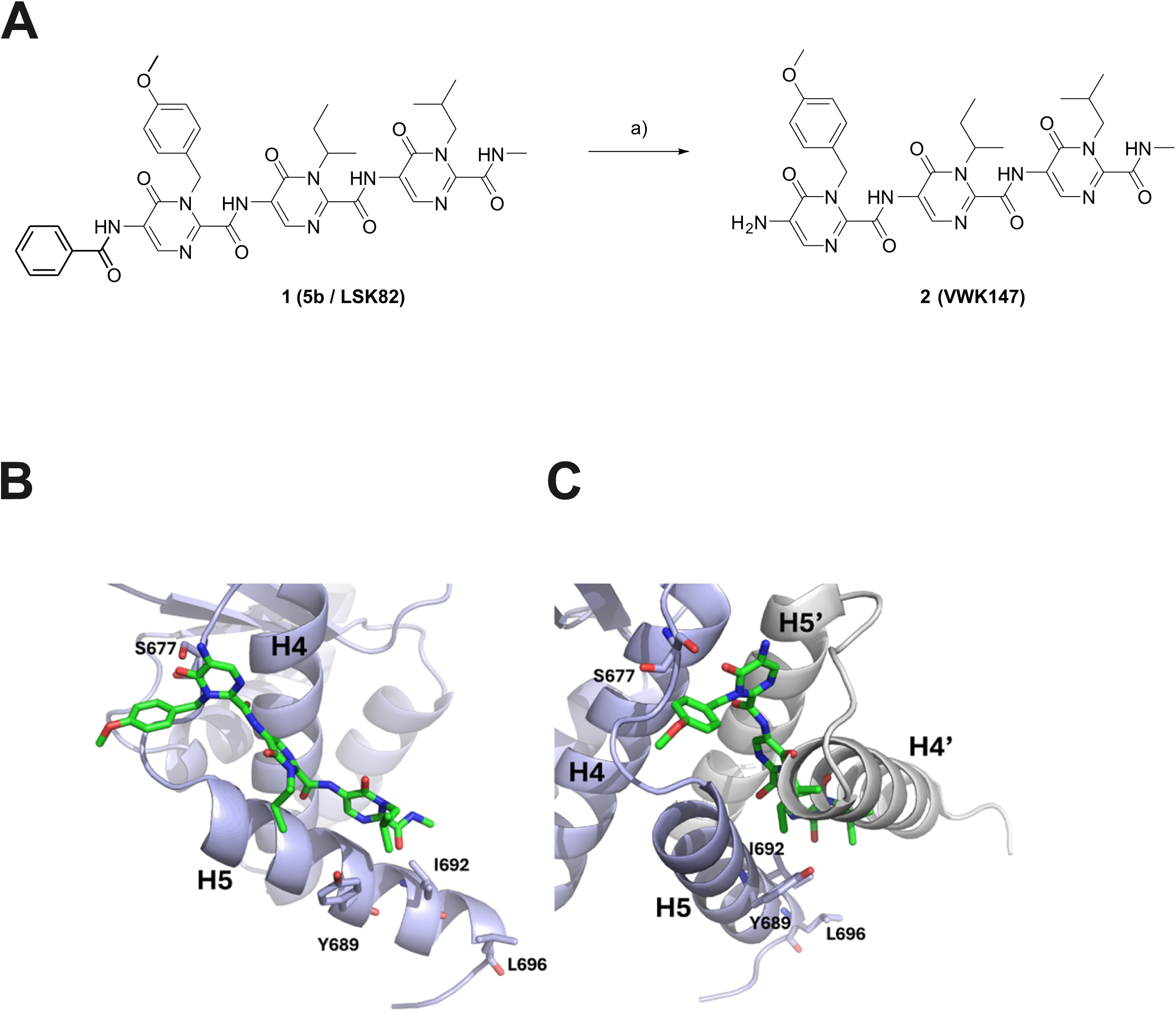
The structure and synthesis of VWK147, and its predicited binding mode at the CTD dimerization interface of an hHSP90 monomer. (A) Synthesis of tripyrimidonamide 2 (VWK147) from benzoyl-protected tripyrimidonamide 1 (LSK82): (a) NaOH, MeOH, 80 °C, 6 h, 40% yield. (B) 2 (VWK147) (green) is predicted to bind to the dimerization interface parallel to helix H5 of the hHSP90 protomer (light blue). The side chains of 2 (VWK147) are in the same order as those it mimics in helix H5’ of the other protomer although the binding mode is slightly shifted towards the N-terminus of H5. There, it forms a hydrogen bond to S677 with its carbonyl group. (C) Upon projection of the binding mode onto the hHSP90 CTD dimer from PDB ID 3Q6M, it becomes evident that the binding mode of 2 (VWK147) effectively precludes H5’ and, notably, H4’ of the opposing protomer (depicted in gray) from engaging with the dimerization interface constituted by H4 and H5 of the initial protomer.

### VWK147 is predicted to bind to the dimerization interface similar to 5b / LSK82

The binding mode of VWK147 at the dimerization interface of the HSP90 CTD was predicted by molecular docking using multiple X-ray structures of a monomeric hHSP90 protomer as a target. The docking results were generally similar in their orientation although the conformation of helix H5 differed between the X-ray structures. They revealed that VWK147 interacts with helices H4 and H5 of the CTD in a manner that would prevent CTD dimerization (Figure 1B and C). In the predicted binding mode, the side chains of VWK147 that mimic residues in helix H5’ of the other protomer are oriented accordingly although the binding pose is slightly shifted towards the N-terminal end of H5 creating a large contact surface with H5. In comparison, the binding mode of 5b / LSK82 predicted previously is flipped, leading to less contact with helix H5 (9). The difference between the two binding modes is likely a result of the missing benzoyl moiety in VWK147. As the conformation of the helices H4 and H5 could change in the absence of the other protomer, we also docked VWK147 to the representative structure of the largest cluster taken from 12 µs of accumulated MD simulations of full-length hHSP90. Here, we predominantly identify predicted binding modes in the cleft between the CTD and the middle domain, as were identified previously for 5b / LSK82 (9). Taken together, the predicted binding modes of VWK147 are similar to those of 5b / LSK82 but VWK147 can form more interactions with H5 at the CTD dimerization interface.

### VWK147 is specific against HSP90 CTD and its co-chaperone function

To investigate the specificity and selectivity of VWK147 against the CTD of HSP90, biochemical cell-free and cellular assays were performed (9). First, we performed a cell-free thermal shift assay to evaluate the binding of VWK147 to recombinant HSP90α CTD. Incubation with VWK147 specifically destabilizes the dimeric HSP90α CTD (ΔT_m_: -8.43 ± 2.75 °C) as indicated by a negative thermal shift (Figure 2A). The thermostabilizing effect was also assessed in a cellular thermal shift assay (CETSA), which is based on ligand-induced stabilization of the target protein in living cells. The quantification of HSP90 was done using capillary-based western blotting and the treatment with VWK147 resulted in a higher amount of ligand-protected intracellular HSP90 with increasing temperature (Figure 2B). Quantification of HSP90 degradation indicated a positive thermal shift (ΔT_m_: 1.66 °C). To further investigate the specificity of VWK147 to the CTD of HSP90, a time-resolved fluorescence resonance energy transfer (TR-FRET) assay was performed. VWK147 disrupts the binding of Peptidyl prolyl isomerase D (PPID, CTD-specific co-chaperone) to the CTD of recombinant HSP90β similar to the C-terminal reference inhibitor coumermycin A1 (CA1) (Figure 2C). To rule out interactions of VWK147 with the HSP90 NTD, a fluorescence polarization (FP) competitive assay was performed using FITC-labelled HSP90 NTD targeting reference inhibitor Geldanamycin (GM). Compared to HSP90-NTD targeting control inhibitors such as PU-H71, Tanespimycin (TM) and GM, VWK147 did not interact with the NTD of HSP90. This observation is consistent with other HSP90 CTD targeting reference inhibitors, such as CA1 and novobiocin (Figure 2D). Next, we investigated the ability of VWK147 to inhibit HSP90 chaperone function in a cell-free luciferase refolding assay. Rabbit reticulocyte lysate was used as a source of HSP90, and it was demonstrated that VWK147 inhibited the recovered luciferase activity in a dose-dependent manner, displaying a comparable efficacy to the established HSP90 NTD targeting inhibitors TM, PU-H71, and AUY922 (Figure 2E). To investigate the effect of VWK147 on the dissociation of the HSP90α CTD, a cell-free assay was conducted using the amine-reactive cross-linker BS^3^ to capture the oligomeric states of HSP90. Increasing concentrations of VWK147 resulted in a reduction of HSP90 CTD dimers, thereby inhibiting the function of HSP90, which is dependent on dimerization (Figure 2F). Collectively, we confirmed that the specificity and selectivity of VWK147 against the CTD of HSP90 and its co-chaperone function are comparable to its predecessor, LSK82.

**Figure 2:**
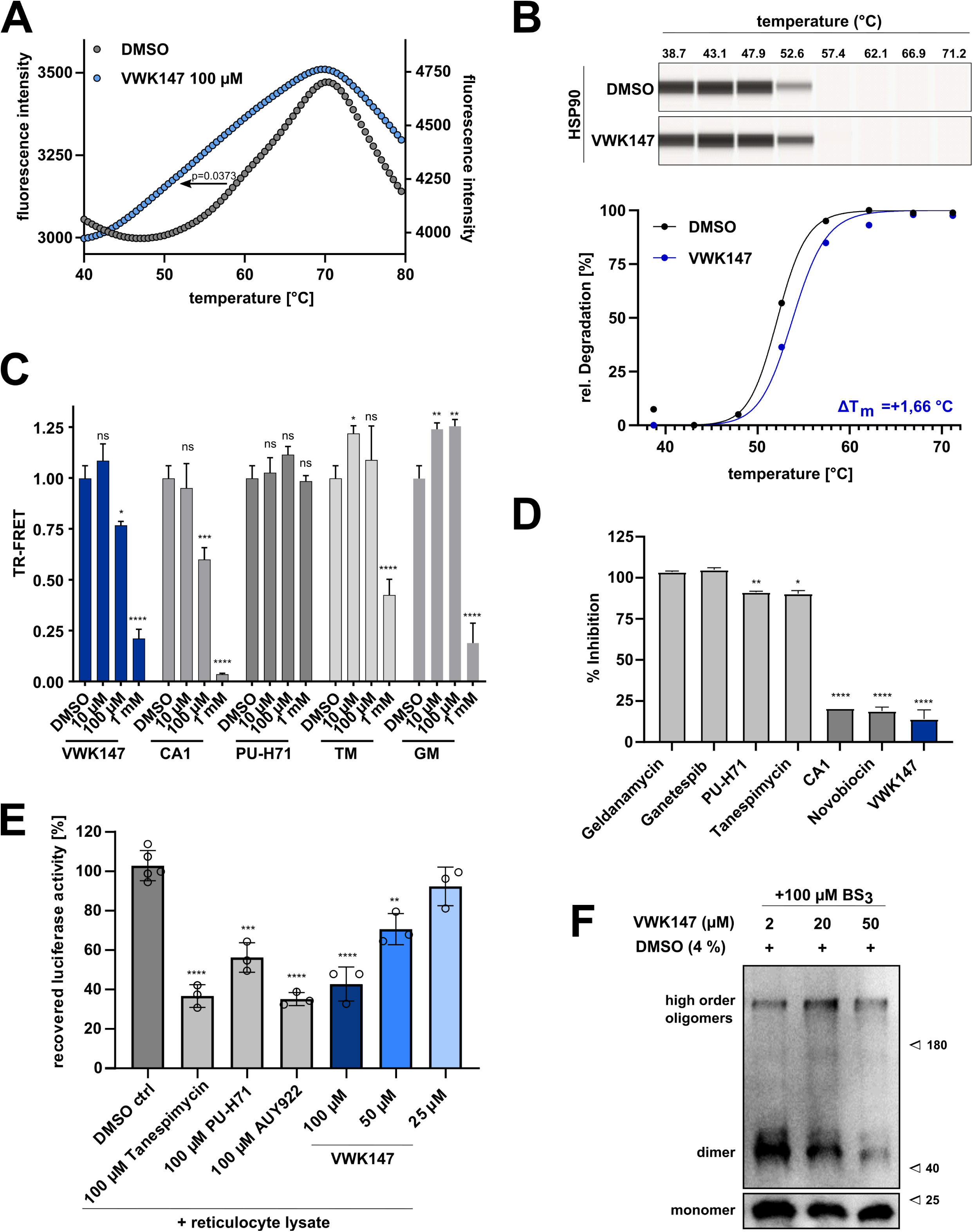
Specificity of VWK147 against HSP90 CTD and its co-chaperone function. **(A)** A cell-free thermal shift assay using recombinant HSP90α CTD protein with VWK147 displays destabilization of HSP90-CTD, indicated by a negative shift. **(B)** Cellular thermal shift assay (CETSA) in K562 leukemia cells displays stabilization of HSP90, indicated by a positive shift. **(C)** VWK147 (1 mM) inhibits HSP90β, comparable to coumermycin A1 (CA1), indicated by a C-terminal FRET assay. **(D)** VWK147 (10 µM) is not binding to the N-terminal domain of HSP90, indicated by an N-terminal fluorescence polarization (FP) assay. (**E**) VWK147 inhibits the HSP90α chaperone function, comparable to Tanespimycin (TM), PU-H71 and AUY922, in the cell-free luciferase-refolding assay, where the incubation of the inhibitors prevented the HSP90- assisted refolding of denatured luciferase. (**F**) Recombinant HSP90α CTD was incubated with BS3 cross-linker at the indicated concentration with VWK147, followed by immunoblotting with the anti-HSP90 (AC88) antibody. VWK147 leads to a reduction of HSP90 dimers. Results are shown as means ± SD of three independent experiments. P values were either determined by a two-tailed unpaired t-test or two-way ANOVA. *p ≤ 0.05; **p ≤ 0.01; ***p ≤ 0.001; ****p ≤ 0.0001.

### VWK147 destabilizes HSP90 client proteins in urothelial carcinoma cells but does not induce a heat shock response

Next, we aimed to confirm VWK147-mediated HSP90 inhibition in urothelial carcinoma cell (UCC) model systems. For that, we employed cisplatin-sensitive and -resistant sublines of the UCCs 253J and T24, respectively (Figure S1). We treated the UCCs with the N-terminal HSP90 inhibitor 17-AAG or VWK147 and monitored protein levels of the HSP90 client proteins ULK1 (29), RIPK1 (30), and CDK4 (31), respectively. ULK1 is an autophagy-inducing kinase, RIPK1 mediates signaling downstream of the TNF-α receptor, and CDK4 regulates cellular transition into S-phase during cell cycle. The levels of the client proteins were reduced upon treatment with either HSP90 inhibitor (Figure 3A-B and Figure S2A-B). We also monitored the induction of a potential heat shock response upon treatment through the detection of HSP27, HSP40 and HSP70 levels by immunoblotting. 17-AAG significantly increased the levels of these heat shock proteins, whereas this was not the case for VWK147 (Figure 3C-D and Figure S2C-D). These data suggest that VWK147 destabilizes HSP90 client proteins without the concomitant induction of heat shock response.

**Figure 3:**
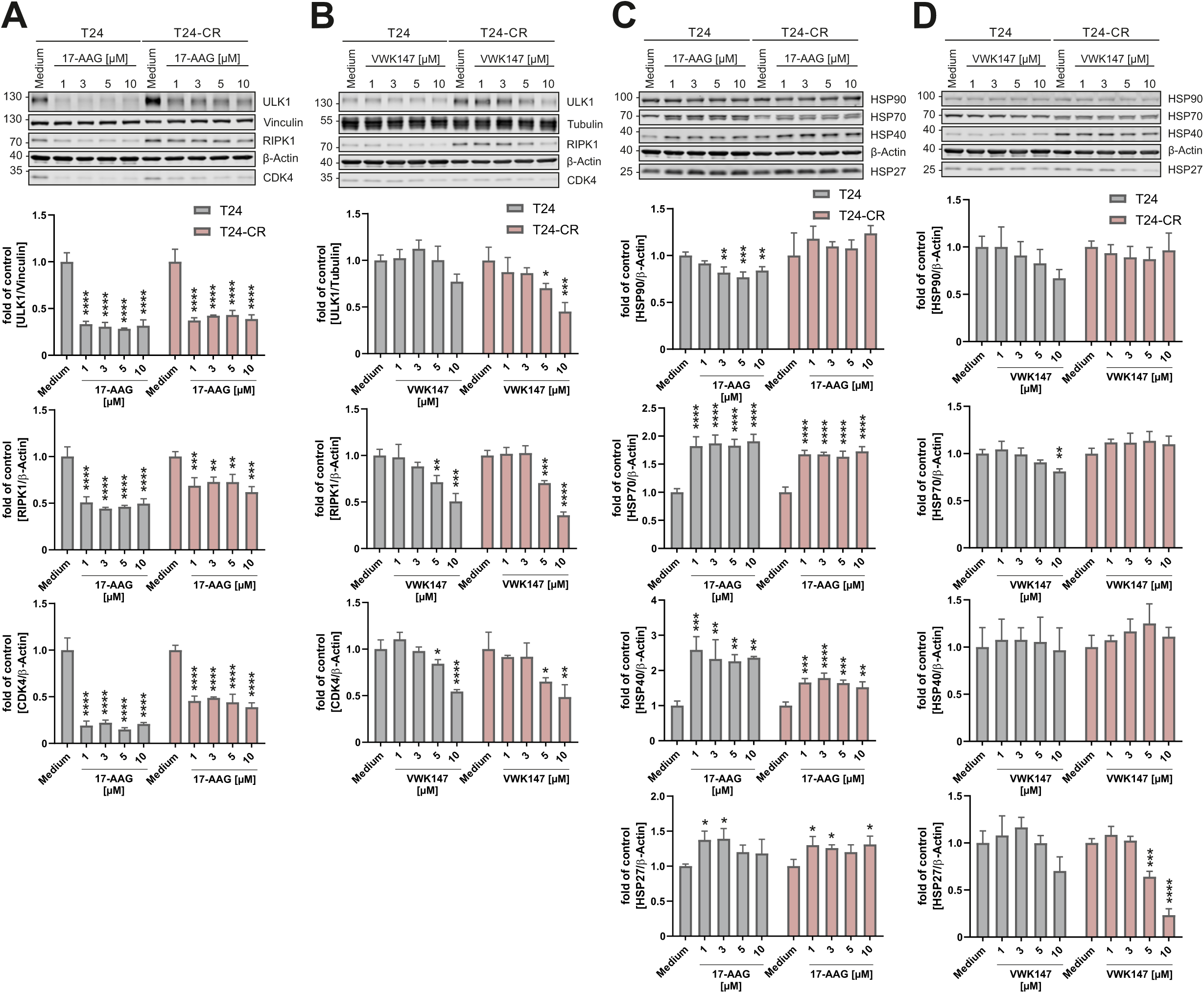
VWK147 destabilizes HSP90 clients but does not induce a heat-shock response. (**A-D**) T24 and T24-CR urothelial carcinoma cells were treated with the indicated concentrations of 17-AAG or VWK147 for 6 h. After treatment, the cells were lysed, and cellular lysates were immunoblotted for ULK1, RIPK1, CDK4, vinculin, tubulin, β-Actin, HSP90, HSP70, HSP40, and HSP27. One representative immunoblot is shown. The quantifications of indicated ratios are from three independent experiments (means + SD). P values were determined by ordinary one-way ANOVA with Dunnett’s multiple comparisons test (comparison to the solvent control of the respective cell line). *p ≤ 0.05; **p ≤ 0.01; ***p ≤ 0.001; ****p ≤ 0.0001.

### VWK147 induces apoptosis in UCCs

Next, we investigated the effect of VWK147-mediated HSP90 inhibition on cell viability in our UCC model systems. Both 17-AAG and VWK147 reduced cell viability in UCCs. Of note, we consistently observed a lower cell viability in the cisplatin-resistant sublines when compared to sensitive sublines, which was evident for both inhibitors (Figure 4A). Both the VWK147-mediated reduction of cell viability and the increased cell death were reduced upon caspase inhibition with QVD, indicating that VWK147—at least partially—induces apoptosis (Figure 4B and 4C). The pro-apoptotic effects of VWK147 were confirmed by the concentration-(Figure 4D) and time-dependent (Figure 4E) detection of cleaved PARP, and by monitoring caspase-3 activation (Figure 4F). Taken together, VWK147 induces apoptosis in cisplatin-sensitive and - resistant UCCs. Of note, cisplatin-resistant cells show higher sensitivity towards VWK147.

**Figure 4:**
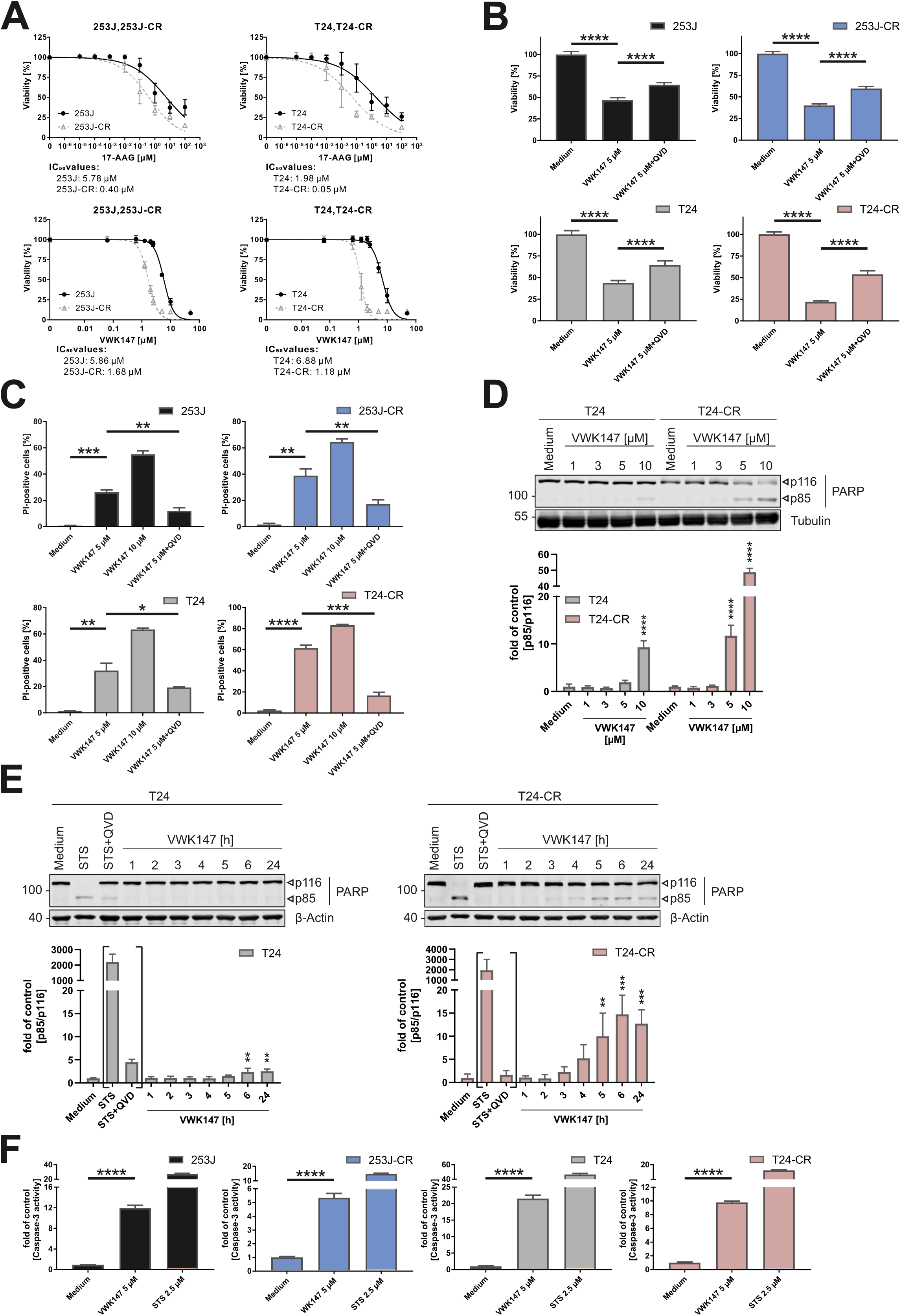
VWK147 reduces cell viability in cisplatin-sensitive and -resistant bladder carcinoma cells, partly by the induction of apoptosis. (**A**) 253J, 253J-CR, T24 and T24-CR urothelial carcinoma cells were treated with indicated concentrations of 17-AAG or VWK147 for 72 h. After treatment, cell viability was measured using Alamar Blue assay. Results are shown as means ± SD of n ≥ 3 independent experiments performed in triplicates for each treatment. (**B**) 253J, 253J-CR, T24 and T24-CR cells were treated with 5 µM VWK147 ± 20 µM QVD (caspase inhibitor) for 24 h. After treatment, cell viability was measured using MTT assay. Results are shown as means + SD of three independent experiments performed in triplicates for each treatment. P values were determined by ordinary one-way ANOVA with Tukeýs post hoc test. ****p ≤ 0.0001 (**C**) 253J, 253J-CR, T24 and T24-CR cells were treated with 5 µM VWK147, 10 µM VWK147 or 5 µM VWK147 + 20 µM QVD for 24 h. After treatment, the cells were stained with propidium iodide and analyzed by flow cytometry. Results are shown as means + SD of three independent experiments performed in triplicates for each treatment. P values were determined by ordinary one-way ANOVA with Tukeýs post hoc test. *p ≤ 0.05; **p ≤ 0.01; ***p ≤ 0.001; ****p ≤ 0.0001 (**D**) T24 and T24-CR urothelial carcinoma cells were treated with the indicated concentrations of VWK147 for 6 h. After treatment, the cells were lysed, and cellular lysates were immunoblotted for PARP and tubulin. One representative immunoblot is shown. The quantifications of cleaved PARP are from three independent experiments (means + SD). P values were determined by ordinary one-way ANOVA with Dunnett’s multiple comparisons test (comparison to the solvent control of the respective cell line). ****p ≤ 0.0001. (**E**) T24 and T24-CR urothelial carcinoma cells were treated with 5 µM VWK147 for the indicated periods of time or with 2.5 µM staurosporine (STS) ± 10 µM QVD for 24 h. After treatment, the cells were lysed, and cellular lysates were immunoblotted for PARP and β-Actin. One representative immunoblot is shown. The quantifications of cleaved PARP are from three independent experiments (means + SD). P values were determined by ordinary one-way ANOVA with Dunnett’s multiple comparisons test (comparison to the solvent control of the respective cell line). Brackets indicate the exclusion of STS and STS+QVD of the significance test. **p ≤ 0.01; ***p ≤ 0.001. (**F**) 253J, 253J-CR, T24 and T24-CR cells were treated with 5 µM VWK147 or 2.5 µM STS for 9 h. Subsequently, DEVDase activity as a surrogate marker for caspase-3 activation was determined via measurement of the fluorescence of the profluorescent caspase-3 substrate DEVD-AMC. Results are shown as means + SD of three independent experiments performed in quadruplicates for each treatment. P values were determined by ordinary one-way ANOVA with Dunnett’s post hoc test. ****p ≤ 0.0001.

### VWK147 sensitizes resistant cells to cisplatin treatment and acts synergistically with mTOR inhibition

Next, we tested whether VWK147 sensitizes the resistant sublines of 253J and T24 towards cisplatin treatment. For this, we treated both sublines either with cisplatin or VWK147 alone or in combination, and monitored PARP and caspase-3 cleavage by immunoblotting. We observed no differences between cells treated with cisplatin alone or with cisplatin + VWK147 in the sensitive subline, whereas in the resistant subline, VWK147 treatment significantly increased cisplatin-induced PARP and caspase-3 cleavage (Figure 5A and 5B). Accordingly, we assume that VWK147 sensitizes resistant cancer cells for cisplatin treatment. The PI3K-AKT-mTOR pathway is a central target for anti-cancer therapy. Several mTOR inhibitors have been assessed in clinical trials. However, these stand-alone approaches revealed limited efficacy due to several reasons, e.g. induction of feedback survival loops, importance of mTOR signaling for healthy tissue, or the induction of autophagy (32). Inhibitors of mTOR were also assessed in several combinatorial therapy approaches, and it has been suggested that inhibition of mTOR can attenuate the HSF1-driven heat shock response (33). Based on this, we decided to analyze whether co-inhibition of mTOR and HSP90 leads to an increase in the cytotoxicity observed in VWK147-treated cisplatin-resistant cells. First, we tested the mTOR inhibitor Torin2 in the UCCs and confirmed its efficacy by detecting reduced phosphorylation of mTOR Ser2448 and the mTOR substrate site Ser758 in ULK1 (Figure S3A and S3B). ULK1 Ser758 is an inhibitory site, and, accordingly, Torin2 treatment resulted in increased phosphorylation of the ULK1 substrate ATG14 at Ser29 (Figure S3A and S3B). To obtain IC_50_ values for combination analyses, we treated cisplatin-sensitive and -resistant 253J and T24 cells with different concentrations of Torin2. The inhibitor efficiently reduced cell viability, with low IC_50_ values especially in the cisplatin-resistant sublines (Figure S3C). To investigate whether Torin2 and HSP90 co-inhibition exhibits a synergistic effect on cytotoxicity, we performed isobologram analysis using the Chou-Talalay method (34). The combination of Torin2 and 17-AAG was not synergistic (Figure 5C), whereas this was clearly the case for VWK147 at most tested concentrations in all four UCC lines (Figure 5D). Collectively, mTOR inhibition and VWK147 treatment synergistically kill UCCs.

**Figure 5:**
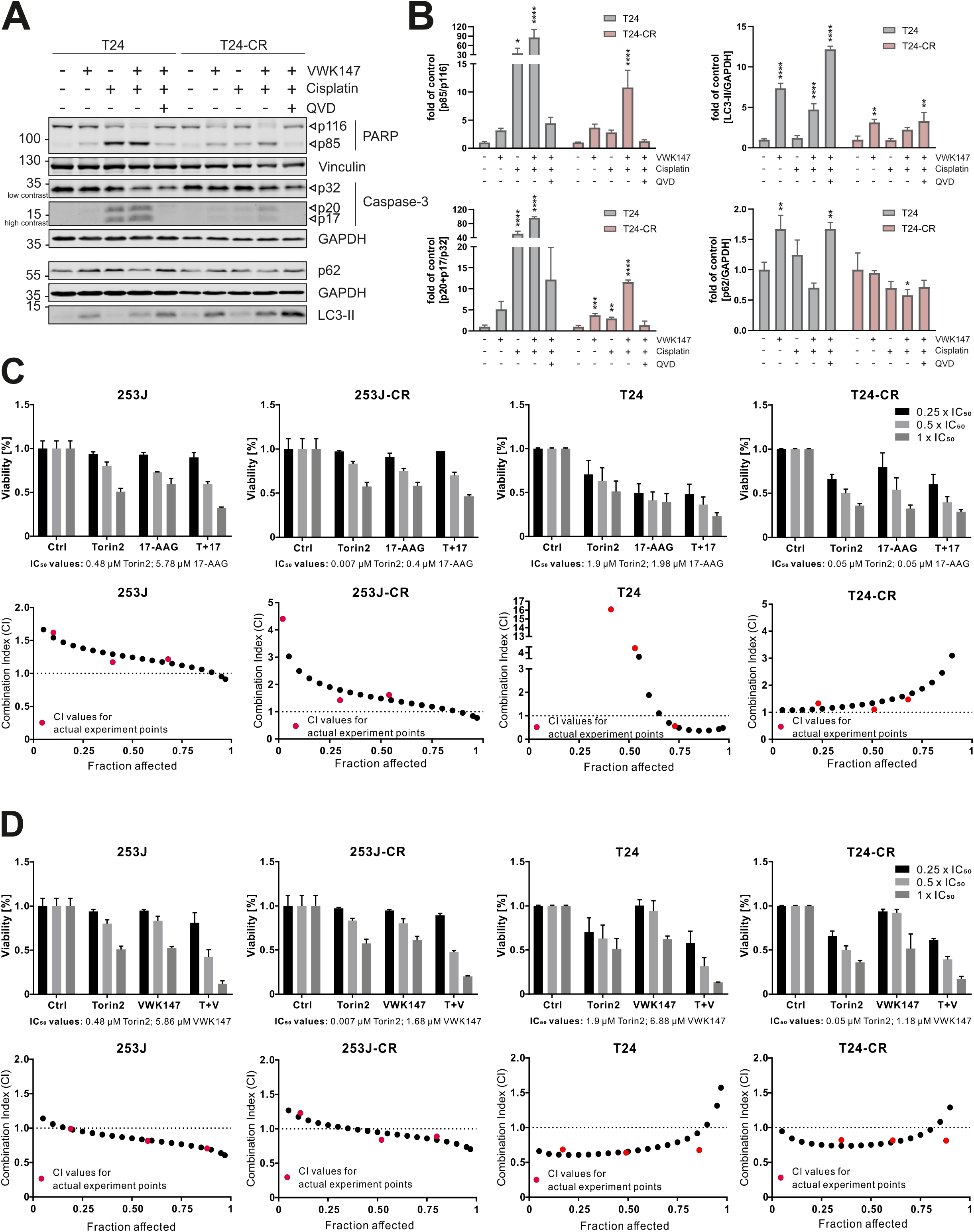
VWK147 sensitizes T24-CR cells to cisplatin-induced apoptosis and acts synergistically with mTOR inhibition. (**A**) T24 and T24-CR urothelial carcinoma cells were treated with IC_50_ concentrations of VWK147 (∼ 10 µM and 5 µM, respectively) and cisplatin (∼ 8 µg/ml and 87 µg/ml, respectively) alone or in combination in absence or presence of 10 µM QVD. After 24 h, the cells were lysed, and cellular lysates were immunoblotted for PARP, Vinculin, Caspase-3, p62, LC3 and GAPDH. One representative immunoblot is shown. (**B**) The quantifications of cleaved PARP, cleaved caspase-3, LC3-II or p62 are from three independent experiments (means + SD). P values were determined by ordinary one-way ANOVA with Dunnett’s multiple comparisons test (comparison to the solvent control of the respective cell line). *p ≤ 0.05; **p ≤ 0.01; ***p ≤ 0.001; ****p ≤ 0.0001. (**C** and **D**) 253J, 253J-CR, T24 and T24-CR cells were treated with Torin2, 17-AAG, VWK147, or Torin2 combined with VWK147 or 17-AAG (0.25 ×, 0.5 × and 1 × IC_50_) for 72 h. The IC_50_ values of each compound were calculated based on the previous mono-treatment. The cell viability was measured using Alamar Blue assay. The combination index values were calculated using CompuSyn (34). The actual experiment points were indicated as red dots whereas simulation dots processed by CompuSyn were indicated as black dots. The combination effects of Torin2 and 17-AAG or VWK147 were determined synergistic (CI < 1), additive (CI = 1) and antagonistic (CI > 1). Values are expressed as means + SD (n = 3).

### Forced expression of HSP70 reduces cytotoxic effect of VWK147

Since we observed the induction of a heat shock response and increased HSP70 expression upon 17-AAG treatment, we hypothesized that this might explain the lack of synergism for Torin2 and 17-AAG. To test this, we exogenously overexpressed 3xFLAG-HSP70 in all four UCC sublines (Figure S4A) and assessed cell viability upon 17-AAG or VWK147 treatment. Indeed, HSP70 overexpression counteracted the reduction in cell viability observed upon HSP90 inhibition in UCCs (Figure S4B). One downstream target of HSP70 is mTOR complex 2 (mTORC2), which mediates the phosphorylation of AKT at Ser473. We observed that 17-AAG increased AKT Ser473 phosphorylation—presumably via the HSP70-mTORC2 axis— whereas this was not the case for VWK147, again supporting the lack of a heat shock response (Figure S4C). Generally, total levels of the HSP90 client kinase AKT were reduced by either HSP90 inhibitor, but the effect on AKT Ser473 phosphorylation was still obvious. It should be noted that mTORC1 activity remained unaffected under all concentrations used for 17-AAG and VWK147, as detected by the phosphorylation status of the mTORC1 substrate p70/S6K at Thr389 (Figure S4D). Next, we assessed AKT Ser473 phosphorylation upon VWK147 treatment in HSP70-overexpressing and vector control cells described above. HSP70 overexpression increased AKT Ser473 phosphorylation, but this was not further modified by VWK147 (Figure S4E). Collectively, our results suggest that the synergism between VWK147 and Torin2 can be at least partially caused by the absence of a heat shock response and therefore a lack of mTORC2 activation and AKT Ser473 phosphorylation. However, we think that the HSP70-mTORC2 axis also plays a minor role for the 17-AAG/Torin2 combination, since Torin2 completely abolishes AKT Ser473 phosphorylation, independent of the presence or absence of 17-AAG (Figure S4F). Thus, other cellular pathways apparently contribute to the VWK147/Torin2 synergism.

### VWK147 induces LC3-II accumulation and inhibits autophagosome-lysosome fusion

Although the lack of a heat shock response upon treatment with VWK147 seems to be relevant for its synergism with Torin2, additional pathways affected by both VWK147 and mTOR inhibition might contribute to this phenomenon. As stated above, mTOR inhibition generally results in the activation of autophagy. In order to investigate whether VWK147 affects autophagy signaling, we analyzed autophagic flux by treating the T24 sublines with the HSP90 inhibitors in the presence or absence of bafilomycin A_1_ (BafA_1_) (Figure 6A). BafA_1_ is an inhibitor of vacuolar H^+^-ATPases and blocks the lysosomal degradation of autophagosomes. VWK147 mono-treatment increased the levels of lipidated LC3 (LC3-II) in either subline, whereas this was not the case for 17-AAG, indicating that VWK147 affects autophagy signaling. In order to distinguish between autophagy-inducing or -inhibiting properties of VWK147, we compared the BafA_1_-treated cells. The combination of VWK147 and BafA_1_ did not significantly increase LC3-II levels compared to BafA_1_ treatment alone, indicating that VWK147 rather inhibits autophagic flux. We also investigated levels of the autophagy receptor p62/SQSTM1. Here, an accumulation was only evident in BafA_1_-treated cells, and the effects of either HSP90 inhibitor were rather negligible (Figure 6A). VWK147-mediated LC3-II accumulation and unaffected p62 levels were also observed in a concentration-dependent manner (Figure 6B).

**Figure 6:**
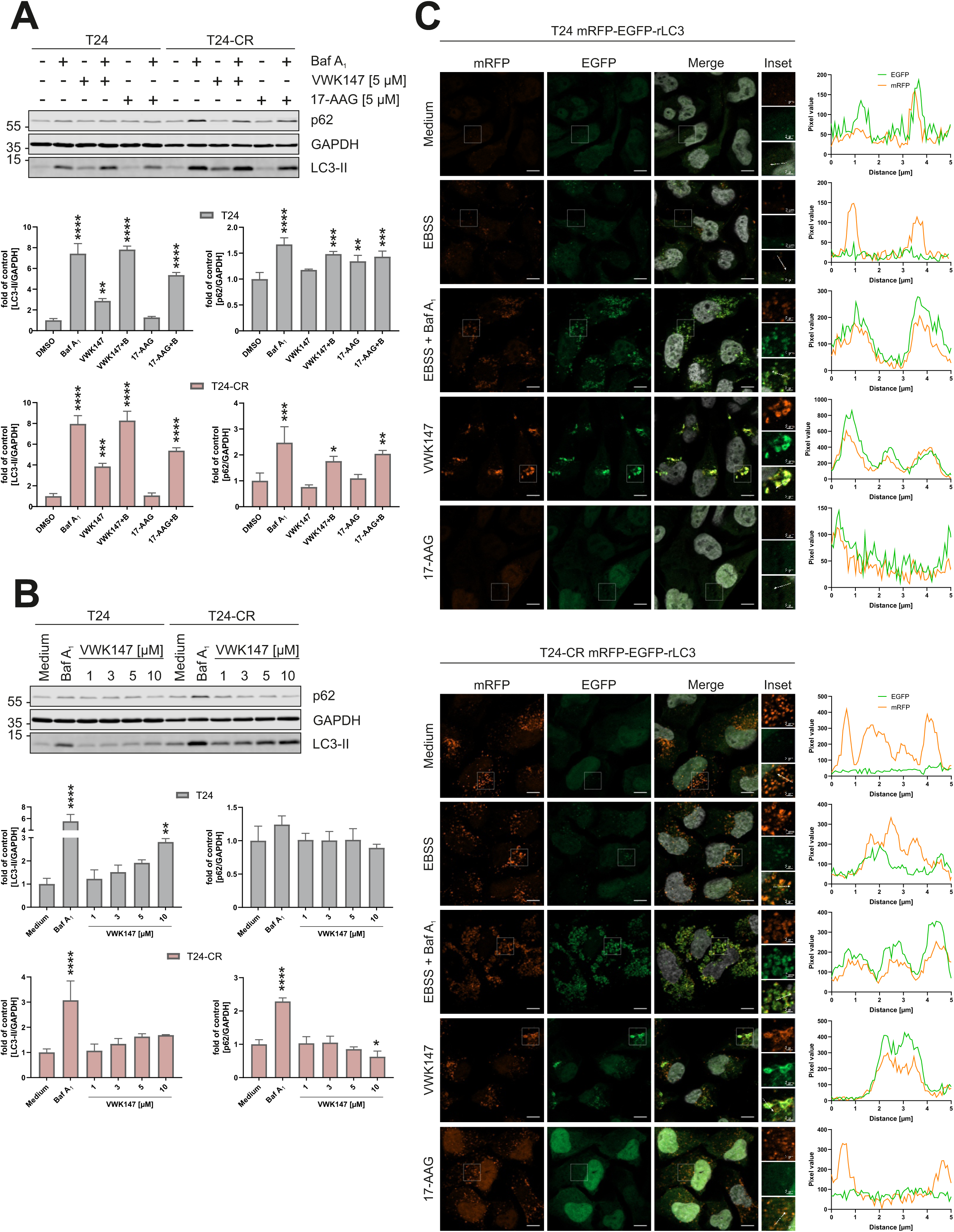
VWK147 inhibits autophagy. (**A**) T24 and T24-CR urothelial carcinoma cells were treated with 5 µM VWK147 or 17-AAG in presence or absence of 20 nM bafilomycin A_1_ for 6 h. After treatment, the cells were lysed, and cellular lysates were immunoblotted for p62, LC3 and GAPDH. One representative immunoblot is shown. The quantifications of indicated ratios are from three independent experiments (means + SD). P values were determined by ordinary one-way ANOVA with Dunnett’s multiple comparisons test (comparison to the solvent control of the respective cell line). *p ≤ 0.05; **p ≤ 0.01; ***p ≤ 0.001; ****p ≤ 0.0001. (**B**) T24 and T24-CR urothelial carcinoma cells were treated with indicated concentrations of VWK147 or 20 nM bafilomycin A_1_ for 6 h. After treatment, the cells were lysed, and cellular lysates were immunoblotted for p62, LC3 and GAPDH. One representative immunoblot is shown. The quantifications of indicated ratios are from three independent experiments (means + SD). P values were determined by ordinary one-way ANOVA with Dunnett’s multiple comparisons test (comparison to the solvent control of the respective cell line). *p ≤ 0.05; **p ≤ 0.01; ****p ≤ 0.0001. (**C**) T24 and T24-CR urothelial carcinoma cells stably expressing mRFP-EGFP-rLC3 were grown on glass coverslips one or two days prior to treatment. Cells were treated with EBSS alone or in combination with 20 nM bafilomycin A_1_, 5 µM VWK147 or 17-AAG for 4 h. Imaging was performed using a Zeiss Axio Observer 7 fluorescence microscope equipped with a 40x/1.4 Oil DIC M27 Plan-Apochromat objective and ApoTome 2. Representative sections are depicted. Scale bars: 10 µM and 2 µM. The line graphs represent the pixel intensities of the areas indicated by the respective dashed white arrows shown in the insets.

In order to ultimately prove the autophagy-inhibiting properties of VWK147, we generated cisplatin-sensitive and -resistant T24 sublines stably expressing mRFP-EGFP-rLC3, and monitored autophagic flux by the detection of yellow puncta (mRFP/EGFP co-localization, representing autophagosomes) and red puncta (mRFP only, representing autolysosomes) using fluorescence microscopy (Figure 6C) (35). As a control for autophagy induction, we used a starvation medium (EBSS). Under starvation conditions, both yellow and red puncta were detectable, suggesting the presence of both autophagosomes and autolysosomes. In contrast, EBSS and BafA_1_ co-treatment abolished red-only dots, confirming the inhibited fusion of autophagosomes with lysosomes. These observations were similar in both cisplatin-sensitive and -resistant sublines, with the exception of rather increased red-only dots in the resistant cells, confirming our previously reported enhanced autophagic flux in cisplatin-resistant T24 (36). Interestingly, VWK147 treatment resulted in an almost complete co-localization of GFP and RFP signals, again indicating that VWK147 interferes with the generation of autolysosomes. Of note, these co-localizing structures appeared to be more tube-like or vesicular aggregates/clusters rather than puncta. In contrast, 17-AAG treatment did not induce similar signals. Collectively, our data show that VWK147 induces LC3 accumulation and inhibits autophagosome-lysosome fusion, and that these effects are not shared by HSP90 inhibitors targeting the NTD.

### VWK147 induces CASM

Since we observed LC3-II accumulation upon VWK147 treatment but rather unaffected p62 levels, we next analyzed if VWK147 induces conjugation of ATG8 proteins to single membranes (CASM). This non-canonical autophagy pathway is independent of the autophagy-inducing ULK1 and PIK3C3 complexes (19). For this, we made use of the PIK3C3 inhibitor SAR405 (37) as control, since it blocks canonical LC3 lipidation. The efficacy of this compound was confirmed in cisplatin-sensitive and –resistant T24, in which it abolished EBSS-induced WIPI2 and LC3 puncta formation (Figure 7A). WIPI2 binds to phosphatidylinositol 3-phosphate, the catalytic product of the PIK3C3 complex, and mediates the recruitment of the ATG8 protein lipidation machinery during canonical autophagy (38). Of note, VWK147-mediated formation of LC3-positive aggregates/clusters was not affected by SAR405 (Figure 7A and Figure S5), pointing towards PIK3C3-independent LC3 lipidation. Although WIPI2 is not required for CASM, we observed that these VWK147-induced LC3-positive structures were frequently also positive for WIPI2, and that especially these double-positive structures were insensitive to SAR405 in both T24 sublines. In order to test for a possible CASM induction by an independent approach, we monitored LC3 lipidation by immunoblotting in different cell lines deficient for different components of the autophagy-inducing ULK1 complex, i.e. *Atg13* knockout murine embryonic fibroblasts, *ATG13/ATG101* double knockout HeLa, and *FIP200* knockout U-2 OS. First, we confirmed the corresponding KOs and an abolished starvation-induced autophagic flux in these cells (Figure S6). VWK147 induced LC3 lipidation in the corresponding wild-type cell lines and also in all tested KO cell lines (Figure 7B-D). Lipidated LC3 levels remained unaffected or were rather reduced by BafA_1_ co-treatment (Figure 7B-D). Furthermore, p62 accumulation was reduced upon VWK147 treatment in a dose-dependent manner in the KO cell lines. In contrast, the N-terminal inhibitor 17-AAG did not evoke a substantial LC3 lipidation or a reduced p62 accumulation in the KO cell lines (Figure S7). Finally, we assessed whether CASM induction might represent a general response to CTD-dependent inhibition of HSP90. For that, we employed novobiocin (39) and confirmed reduced expression of the client proteins AKT and ULK1 (Figure S8A). Similar to VWK147, novobiocin induced LC3 lipidation and reduced p62 levels in our UCC models (Figure S8B and S8C) and in *FIP200* KO U-2 OS (Figure S8D). LC3 lipidation was again partially reduced by concomitant BafA_1_ treatment, suggesting that a BafA_1_-sensitive non-canonical LC3 lipidation might represent a general phenomenon of CTD-dependent HSP90 inhibition. However, it needs to be mentioned that novobiocin concentrations were 200-500 fold higher than VWK147 concentrations.

**Figure 7:**
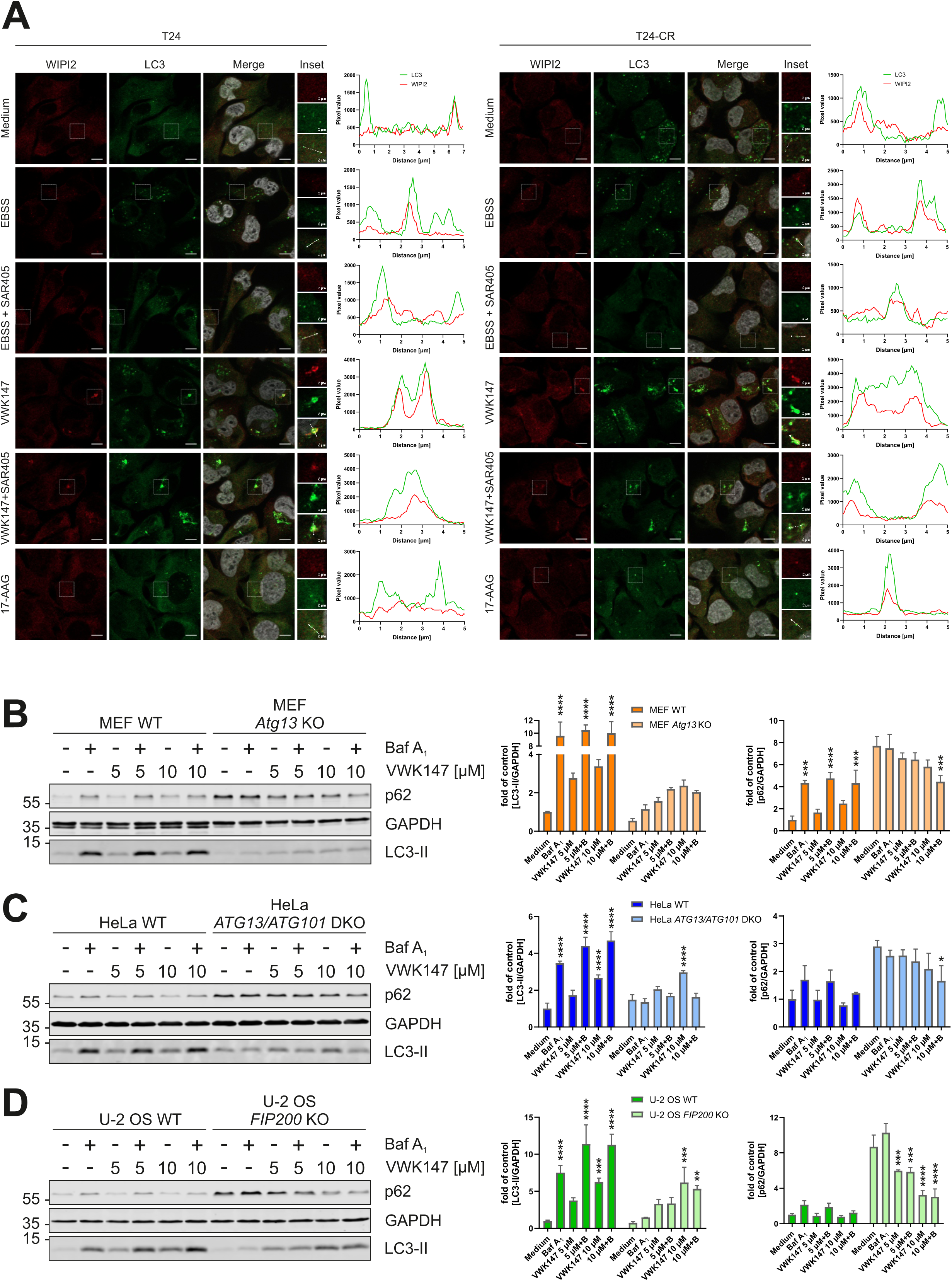
VWK147 induces non-canonical autophagy. (**A**) T24 and T24-CR urothelial carcinoma cells were grown on glass coverslips one or two days prior to treatment. Cells were treated with EBSS or 5 µM VWK147 alone or in combination with 5 µM SAR405, or 5 µM 17-AAG. Imaging was performed using a Zeiss Axio Observer 7 fluorescence microscope equipped with a 40x/1.4 Oil DIC M27 Plan-Apochromat objective and ApoTome 2. Representative sections are depicted. Scale bars: 10 µM and 2 µM. The line graphs represent the pixel intensities of the areas indicated by the respective dashed white arrows shown in the insets. See also Figure S5. (**B**) MEF WT and MEF *Atg13* KO, (**C**) HeLa WT and HeLa *ATG13/ATG101* DKO and (**D**) U-2 OS WT and U-2 OS *FIP200* KO were treated with indicated concentrations of VWK147 in absence or presence of 20 nM bafilomycin A_1_ for 6 h. After treatment, the cells were lysed, and cellular lysates were immunoblotted for p62, LC3 and GAPDH. One representative immunoblot is shown. The quantifications of indicated ratios are from three independent experiments (means + SD). P values were determined by ordinary two-way ANOVA with Tukeýs multiple comparisons test (comparison to the solvent control of the respective cell line). *p ≤ 0.05; **p ≤ 0.01; ***p ≤ 0.001; ****p ≤ 0.0001.

Altogether, our data indicate that VWK147 1) inhibits HSP90 without inducing a heat shock response, 2) re-sensitizes cisplatin-resistant cancer cells to cisplatin, 3) acts synergistically with mTOR inhibition, 4) inhibits canonical autophagy, and 5) concomitantly induces CASM-like processes.

## Discussion

Targeting HSP90 has become a promising approach in anticancer therapy. Several cancer-associated proteins are regulated by HSP90, and various HSP90 inhibitors have been assessed in clinical trials (1). Frequently, therapeutic interventions are antagonized by cellular stress responses. In the case of HSP90 inhibition, its efficacy is hampered by the induction of HSR. Additionally, autophagy is also employed by cancer cells in order to adapt to metabolic stress and/or chemotherapy. Here, we reported the synthesis, biochemical characterization and cell biological assessment of a small molecule inhibitor of C-terminal HSP90 dimerization that additionally interferes with two central survival pathways in cancer cells, i.e. this compound does not induce HSR and blocks canonical autophagy.

In recent years, the crosstalk between HSP90 and autophagy has been investigated, and several autophagy-regulating proteins have been identified as HSP90 client proteins (40). HSP90 has been reported to regulate Toll-like receptor (TLR)-induced autophagy by interacting with Beclin 1, which is a core component of the autophagy-inducing VPS34/PIK3C3 complex (41). Similarly, ULK1-mediated mitophagy is regulated by the HSP90-Cdc37 chaperone complex (29). Also chaperone-mediated autophagy (CMA) is regulated by HSP90, as lysosome-associated HSP90 ensures the stability of LAMP-2A at the lysosomal membrane (42). Accordingly, it is not surprising that HSP90 inhibition affects autophagy signaling. Of note, both autophagy-inducing and -inhibiting properties of HSP90 inhibitors have been reported. Generally, HSP90 inhibition-mediated induction of autophagy might be explained by the inactivation of client proteins such as AKT or mTOR. At the same time, these pro-autophagic effects might be counteracted by the inactivation of pro-autophagic client proteins such as ULK1. Additionally, different cellular model systems and/or differential effects on starvation-induced vs. basal autophagy have to be considered when analyzing autophagy modulation by HSP90 inhibition. Joo et al. reported that 17-AAG inhibits starvation-induced autophagy (29). Impaired autophagic flux was also observed in A549 lung cancer cells exposed to coumarin pyrazoline compounds that might represent HSP90 inhibitors (43). Mechanistically, it has also been suggested that HSP90 inhibition diminishes the expression of ATG7, thereby impeding autophagic responses (44). In our case, this effect does not appear to play a major role, since we observe strong LC3 lipidation upon VWK147 mono-treatment. In contrast to the anti-autophagic effects listed above, several reports describe the induction of autophagy upon HSP90 inhibition (45–48). Schaefer et al. observed that co-treatment consisting of 17-AAG and the kinase inhibitor D11 induces autophagy (46). D11 causes the suppression of the heat shock response, and thus 17-AAG/D11 co-treatment might be mimicked by our single treatment with VWK147. However, our analyses using the RFP-EGFP-rLC3 reporter construct indicate an inhibition of autophagy upon VWK147 treatment. This VWK147-mediated inhibition of autophagy may contribute to its cytotoxicity in our UCC models. We have previously shown that autophagy-regulating proteins are up-regulated in cisplatin-resistant UCCs (36), possibly indicating an autophagy-addicted state of these cells. Along these lines, cisplatin-resistant cells were more sensitive to VWK147 than the cisplatin-sensitive subline. Accordingly, HSP90 inhibition and inhibition of late stage autophagy are molecular properties that are combined in VWK147 and that might be central for its therapeutic efficacy.

Non-canonical LC3 lipidation represents an autophagy-related signaling pathway that gained recent interest (19). The conjugation of ATG8 proteins to single membranes (CASM) occurs independently of ULK1- and VPS34/PIK3C3-complexes, and this is also the case for VWK147-induced LC3 lipidation, as shown by the usage of the VPS34/PIK3C3 inhibitor SAR405 or different cell lines deficient for different components of the ULK1 complex. Interestingly, we also observe VPS34/PIK3C3-dependent phenomena upon VWK147 treatment, such as formation of WIPI2-positive aggregates/clusters. It has been reported that starvation-induced WIPI2 puncta formation is inhibited by Wortmannin, but that the association of WIPI2 with membranes remains unaffected (49). The VWK147-induced LC3-positive aggregates/clusters are clearly different from starvation-induced puncta and frequently reveal a vesicular or tube-like morphology. These structures might be membraneous, and thus WIPI2 association is possible. Generally, it appears that VWK147 blocks canonical autophagy in cells that are autophagy-competent (i.e. harboring the autophagy-mediating machinery), whereas VWK147-mediated non-canonical LC3 lipidation is clearly detectable in cells lacking the components of the ULK1 complex. Accordingly, the LC3/WIPI2-positive clusters observed in T24 cells may also represent stalled autophagosomes and not necessarily CASM-related structures. It should also be noted that CASM is generally dependent on the vacuolar H^+^-ATPase and thus blocked by BafA_1_. Partially we observe this dependency, but sometimes VWK147-induced LC3-II levels remain unaffected by BafA_1_ also in cells lacking the components of the ULK1 complex. Interestingly, both VWK147 and novobiocin reduced p62 levels in wild-type cells and in cells deficient for components of the ULK1 complex, and especially in the latter this reduced p62 accumulation was not sensitive to BafA_1_ treatment. Taken together, the relative contributions of 1) the inhibition of canonical autophagy and 2) the induction of non-canonical LC3 lipidation need to be characterized in future studies.

We also observe a clear difference between 17-AAG and VWK147 on ULK1 levels. It has previously been described that 17-AAG treatment results in a faster-migrating, presumably hypophosphorylated form of ULK1 (29). In contrast, VWK147 treatment does not alter the migrational behavior of ULK1 but reduces its overall levels. These differential observations might point towards differential effects of these two inhibitors on the “client spectrum” of upstream kinases phosphorylating ULK1. Similarly, we do not observe non-canonical LC3 lipidation for 17-AAG, whereas this is the case for the CTD-targeting inhibitors VWK147 and novobiocin. Collectively, differential effects of NTD- and CTD-targeting HSP90 inhibitors await further clarification.

Söti et al. reported that cisplatin represents a selective nucleotide competitor of the C-terminal ATP-binding site in HSP90 (50). Accordingly, simultaneously targeting HSP90 CTD dimerization with VWK147 and nucleotide binding with cisplatin may effectively abolish HSP90 CTD activity. Since HSP90 CTD activity is critical for stabilizing numerous client proteins, this dual inhibition strategy suggests that VWK147 could serve as an effective combination partner with cisplatin.

The PI3K/AKT/mTOR pathway is an essential pathway that regulates various cellular processes, including cell growth, proliferation, survival and autophagy (51, 52). In several cancers, the mTOR signaling pathway is dysregulated (53). Accordingly, mTOR has emerged as an attractive therapeutic target, and both allosteric (rapamycin and analogs, i.e. rapalogs) and ATP-competitive kinase inhibitors (TORKinibs) have been or are currently tested in clinical trials (32). The efficacy of monotherapies targeting mTOR is hampered by the induction of feedback survival loops (32), resulting in the development of combinatorial approaches. There exists a crosstalk between mTOR and HSP90 signaling. On the one hand, mTOR inhibition can help to overcome HSP90-induced heat-shock responses via the suppression of HSF1 and HSP70 expression (33). On the other hand, HSP70 associates with Rictor and is required for mTORC2 formation and activity (54). In recent years, the combination of HSP90 and mTOR inhibitors has been shown to be an effective therapeutic approach in various cancer models (27, 55–59). Recently, Pan et al. reported the synthesis and evaluation of a dual HSP90/mTOR inhibitor (27). The authors describe an *in vivo* anti-tumor activity of their compound in UCC xenograft models. However, efficacy in cisplatin-resistant cells was not assessed and awaits further analysis. Kim et al. reported that the N-terminal HSP90 inhibitor 17-DMAG and the PI3K/mTOR inhibitor NVP-BEZ235 show a synergistic anti-tumor effect in cisplatin-resistant T24 cells (59). We do not observe synergism between the N-terminal inhibitor 17-AAG and Torin2, but this may be due to the different nature of applied inhibitors and/or different cellular model systems. However, we do observe synergism between VWK147 and mTOR inhibition in cisplatin-resistant cells. We propose that this is caused by two VWK147-dependent effects on cellular survival responses: 1) lacking induction of heat-shock response, and 2) inhibition of autophagy. We have previously reported that autophagy-related proteins are up-regulated in different cisplatin-resistant UCCs compared to the sensitive parental cell lines (36), and accordingly, these cells may be particularly susceptible to treatment with autophagy-modulating compounds. The induction of autophagy and the simultaneous blockade of this process have been proposed as anti-cancer strategy (60–62), and this might exactly be the mechanism we observe here: the induction of early autophagy events by Torin2, and the blockade of late stage autophagy by VWK147. Certainly, the relevance and/or contribution of the described non-canonical LC3 lipidation awaits further clarification. Nevertheless, VWK147 represents a promising inhibitor of the C-terminal HSP90 dimerization with double-edged autophagy-modulating properties, and this compound can—as monotherapy or in combination with cisplatin or mTOR inhibitors—prove useful in order to optimize anti-cancer therapies and/or to overcome therapy resistance.

## Acknowledgements

We thank Margaretha Skowron (Department of Urology, Medical Faculty, Heinrich Heine University, Düsseldorf, Germany) for providing cisplatin-sensitive and -resistant 253J and T24 bladder carcinoma cells. We thank Toshio Kitamura (Institute of Medical Science, University of Tokyo, Japan) for providing Plat-E cells, Noboru Mizushima (University of Tokyo, Japan) for providing wild-type and *Atg13* KO MEFs, and Fulvio Reggiori (Aarhus University, Denmark) for providing wild-type and *FIP200* KO U-2 OS cells. We are grateful for computational support and infrastructure provided by the “Zentrum für Informations- und Medientechnologie” (ZIM) at the Heinrich Heine University Düsseldorf and the computing time provided by the John von Neumann Institute for Computing (NIC) on the supercomputer JUWELS at Jülich Supercomputing Centre (JSC) (user ID: VSK33).

## Author contributions

CD and YS performed cell viability assays, immunofluorescence microscopy, immunoblot analyses, and generated the cell lines expressing 3xFLAG-HSP70 or mRFP-EGFP-rLC3. VW synthesized VWK147. CG performed molecular docking and simulations, CG and HG analyzed the modeling results. MV performed thermal shift, luciferase refolding and BS3 crosslinker assays. ND performed CETSA, nFP and c-terminal FRET assays. DS, LB, AF, SA, MJM and SW gave technical support. CD, YS, VW, MV, ND, SB, TK and BS analyzed and interpreted the data. CD, YS, VW, MV, ND, CG, HG, SB, TK and BS wrote the manuscript. SB, TK and BS supervised the project. All authors discussed the results and commented on the manuscript.

## Materials and methods

### Procedure for the synthesis of tripyrimidonamide **2** (VWK147)

A solution of 1.05 g tripyrimidonamide **1** (1.35 mmol) in 6 ml of a 0.5 M methanolic sodium hydroxide was heated to 80 °C. After 5 h the solvent was evaporated using a rotary evaporator. The resulting yellow solid was purified using flash column chromatography (n-hexane:ethyl acetate). The main fractions were combined, and the solvent removed using a rotary evaporator. The bright yellow solid was recrystalized using n-hexane:ethyl acetate to yield **2**.

### Analytical data for tripyrimidonamide **2** (VWK147)

**Yield**: 40%; 362 mg (0.54 mmol), yellow solid

**Mp**: 226 °C (dichloromethane)

**HPLC**: R_t_: 15.57 min, purity: 97.2%

**^1^H NMR** (300 MHz, chloroform-*d*) δ 10.44 (s, 1H), 10.11 (s, 1H), 8.96 (s, 1H), 8.93 (s, 1H), 7.84 – 7.65 (m, 1H), 7.38 (s, 1H), 7.31 (d, *J* = 8.2 Hz, 2H), 6.81 (d, *J* = 8.2 Hz, 2H), 5.98 (s, 2H), 2.06 – 1.86 (m, 1H), 5.31 – 5.11 (m, 2H), 4.66 (d, *J* = 6.7 Hz, 2H), 3.75 (s, 3H), 2.99 (s, 3H), 2.36 – 2.20 (m, 1H), 2.18 – 1.90 (m, 2H), 1.68 (d, *J* = 6.0 Hz, 3H), 0.93 (d, *J* = 6.2 Hz, 6H), 0.84 (t, *J* = 7.1 Hz, 3H).

**^13^C NMR** (75 MHz, chloroform-*d*) δ 160.92, 158.99, 158.89, 158.73, 157.94, 157.88, 144.75, 144.31, 136.83, 136.52, 134.20, 132.99, 129.64, 129.01, 128.29, 126.98, 123.50, 113.85, 59.98, 55.23, 51.03, 46.56, 28.81, 26.65, 25.88, 19.82, 17.87, 11.36.

**HR-MS (ESI+):** calculated for [C_32_H_38_N_10_O_7_+H^+^] m/z: 675.2998; found: 675,2995

### Thermal shift assay

The thermal shift assay was performed as previously described (9). CTD r-HSP90α (5 µM) protein and VWK147 (100 µM) were mixed together in the assay buffer (1× PBS, pH = 7.5) and were incubated for 2 h. Then, 6× SYPRO orange dye (Sigma-Aldrich) was added to the mixture to a final volume of 20 μl. 96-well polymerase chain reaction (PCR) plates and a PCR system (BioRad, CFX Connect real-time system) were used to heat the samples from room temperature to 95 °C in increments of 0.5 °C for 10 s, with the excitation wavelength at 470 nm and emission wavelength at 570 nm. For a determination of protein melting temperature values (Tm), the melting curve for each data set was analyzed by GraphPad Prism 8.0.2 and fitted with the sigmoidal Boltzmann fit. Melting temperatures without the inhibitors (DMSO) were used as a control.

### Cellular thermal shift assay (CETSA)

A CETSA assay was performed as described previously (9). K562 cells were incubated with VWK147 (or DMSO) for 24 h. Cells were harvested by centrifugation (400g for 5 min at room temperature) and washed five times with ice-cold PBS. The pellets were dissolved in PBS and later equally divided into 200 μl PCR tubes. Solutions were heated at the indicated temperature gradient for 3 min and 30 s (T-Gradient Cycler, Biometra). Aliquots were then snap-frozen in liquid nitrogen and thawed at 25 °C in a thermal cycler (GeneAMP PCR System2700, Applied Biosystems) three times, followed by centrifugation at 10 000 g for 20 min at 4 °C. The supernatants were harvested, and protein levels were measured by a quantitative simple western immunoassay (JESS, BioTechne, Minneapolis, MN). JESS was performed following the manufacturer’s instructions. Protein levels represented by the area under the curve of the electropherograms were normalized to the lowest temperature set as 0% degradation. The ΔTm value was determined by plotting normalized data using a sigmoid dose curve and nonlinear regression (GraphPad Prism 8.0.2).

### Time-resolved fluorescence resonance energy transfer (TR-FRET) assay

An evaluation of the C-terminal HSP90 binding affinity to PPID (cyclophilin D) was performed using the HSP90 CTD TR-FRET assay kit (50289, BPS Bioscience, San Diego, CA). For the positive control, the inhibitor was substituted for DMSO, and for the negative control, PPID-GST-tag was substituted for 1x HSP90 assay buffer. The assay was performed as described previously (9). Samples were incubated for 2h at room temperature protected from light and measured with a microplate-reader (SPARK10M, Tecan). Fluorescence was measured using a time-resolved reading mode with two subsequent measurements: The first measurement was performed using a 340 nm/620 nm (excitation/emission) wavelength with a lag time of 60 μs and integration time of 500 μs. The second measurement was performed using a 340 nm/665 nm (excitation/emission) wavelength with a lag time of 60 μs and integration time of 500 μs. Data analysis was performed using the TR-FRET ratio (665 nm emission/620 nm emission). The TR-FRET ratios are normalized to % activity by setting the negative control as 0% activity and the positive control as 100% activity [(FRET_sample_ – FRET_neg_)/(FRET_pos_ – FRET_neg_) * 100%]. Two-way ANOVA was performed to calculate significance against DMSO control.

### Fluorescence polarization (FP) assay

An evaluation of the binding affinity of compounds toward the ATP pocket of HSP90 NTD was determined by a competitive binding assay against FITC-labeled geldanamycin (GM) using the HSP90 NTD assay kit (50293, BPS Bioscience) as described previously (9). The inhibitor sample wells were filled with 1X HSP90 assay buffer, supplemented with DTT, BSA, FITC-labeled GM (100 nM), and 10 μl of VWK147 (final concentration 10 µM). The reaction was initiated by adding 20 μl of Hsp90 (17 ng/μl) and incubating at room temperature for 3 h with slow shaking. Background wells (master mix only), negative controls (FITC-labeled GM, buffer, and DMSO), and positive controls (FITC-labeled GM, buffer, DMSO, and Hsp90) were also included within the assay plate. Fluorescence polarization was measured at a 470 nm excitation wavelength and 525 nm emission wavelength in a microtiter-plate reader (Infinite M1000pro by Tecan). The instrument-specific g-factor of 1.187 was used to calculate the polarization (I_II_ – G(I┴)/ (I_II_ + G(I┴))*1000, whereas the percentage of HSP90-bound FITC-GM was calculated using P_norm_ = (P_Inhibitor_-P_neg_) / (P_pos_ – P_neg_)*100. Statistical analysis was performed utilizing two-way ANOVA against unlabeled Geldanamycin.

### Luciferase refolding assay

A luciferase refolding assay was performed as previously described (9). Recombinant firefly luciferase from Photinus pyralis (Sigma-Aldrich, St. Louis, MO; 10 × 1010 units/mg) was diluted (1:100) in denaturation buffer (25 mM Tricine, pH 7.8, 8 mM MgSO4, 0.1 mM EDTA, 1% Triton X-100, 10% glycerol, and 10 mg/ml BSA) at 38 °C for 8 min. Rabbit reticulocyte lysate (Promega, Madison, WI) was diluted 1:1 by the addition of cold mix buffer (100 mM Tris, pH 7.7, 75 mM Mg(OAc)2, 375 mM KCl, and 15 mM ATP), creatine phosphate (10 mM), and creatine phosphokinase (16 U/ml) and was preincubated at 30 °C with VWK147 and controls (DMSO, Tanespimycin, PU-H71, and AUY922) for 1 h. Afterward, 1 μl of denatured luciferase or active luciferase (as a control) was added to 20 μl of a rabbit reticulocyte mixture. As a control, denatured or active luciferase was incubated without reticulocyte lysate in buffer containing 20 mM Tris, pH 7.5, 150 mM NaCl, 1% hemoglobin, and 4% BSA. At desired time points, 1.5 μl samples were removed and added to 40 μl of assay buffer (25 mM Tricine, pH 7.8, 8 mM MgSO4, 0.1 mM EDTA, 33 μM DTT, 0.5 mM ATP, and 0.5 mM luciferin), and the luminescence was read using a Spark microplate reader (Tecan). Percent luciferase refolding was determined by comparing the samples to DMSO (100 %).

### BS3 crosslinker assay

Hsp90 CTD dimerization was evaluated using an amine-reactive chemical cross-linker bis(sulfosuccinimidyl) suberate (BS3) (Pierce) as previously described (DOI: 10.1021/acscentsci.2c00013). HSP90α CTD protein (2 μM) was diluted in Na_2_HPO_4_ (25 mM; pH 7.4) and treated with different concentrations of the inhibitor VWK147 to make a final volume of 25 μl. The reaction mixture was incubated at room temperature for 1 h. The amine-reactive cross-linker BS3 was added to a final concentration of 100 μM, and the samples were incubated for 1 h at room temperature. Cross-linking was quenched by the addition of SDS sample buffer and subsequent heating for 5 min at 95 °C. Samples were run in 12% SDS-PAGE gels followed by Western blotting. Blots were probed with anti-HSP90 (AC88, Abcam) antibody.

### Reagents

Torin2 (#S2817) and SAR405 (#S7682) were purchased from Selleck Chemicals (Houston, TX, USA); bafilomycin A_1_ (#B1793) was purchased from Sigma-Aldrich (St. Louis, MO, USA). Additionally, the following reagents were used: dimethyl sulfoxide (DMSO; AppliChem GmbH, #A3672), staurosporine (STS; [Biozol, #S-9300]), Q-VD-Oph (QVD; [Selleck Chemicals, #S7311]), milk powder (Carl Roth, #T145.2), PBS (Thermo Fisher Scientific/Gibco, #14190-094), 0.05% trypsin/EDTA solution (Thermo Fisher Scientific/Gibco, #25300-062), hygromycin (InvivoGen, #ant-hg-1), puromycin (InvivoGen, #ant-pr-1) and Ac-DEVD-AMC (Biomol, #ABD-13402). For immunoblotting, primary antibodies against SQSTM1/p62 (PROGEN Biotechnik, #GP62-C), LC3 (Cell Signaling Technology, #2775), GAPDH (abcam, #8245), β-Actin (Sigma-Aldrich, #A5316), phospho-mTOR Ser2448 (Cell Signaling Technology, #2971), mTOR (CST, #2972), phospho-ULK1 Ser757 (CST, #6888), ULK1 (clone D8H5, CST, #8054), ATG14 (MBL, #PD026; CST, #5504), phospho-ATG14 Ser29 (CST, #92340), Vinculin (Sigma-Aldrich, #V9131), tubulin (Sigma, #T5168), poly (ADP-ribose) polymerase (PARP)-1 (Enzo, #MBL-SA250; CST, #9542), Caspase-3 (CST, #9662), RIPK1 (BD Biosciences, #610459), CDK4 (Thermo Fisher Scientific, #MA5-13720), HSP90 (CST, #4875), HSP70 (Abcam, #ab2787), HSP40 (CST, #4871), HSP27 (CST, #2402), phospho-AKT Ser473 (CST, #9271), AKT (CST, #9272), FLAG (Sigma-Aldrich, #F1804), phospho-p70/S6K Thr389 (CST, #9206), p70/S6K (CST, #9202), FIP200 (Proteintech, #17250-1-AP), ATG13 (Sigma-Aldrich, #SAB4200100) and ATG101 (CST, #13492) were used. IRDye 680- or IRDye 800-conjugated secondary antibodies (#926-68077, #926–68072/73, #926-32212/13) were purchased from LI-COR Biosciences.

For immunofluorescence, primary antibodies against WIPI2 (Biorad, #MCA5780GA) and LC3B (MBL, #PM036) were used. Alexa Fluor^®^ 488-conjugated and Alexa Fluor^®^ 647-conjugated secondary antibodies were purchased from Jackson ImmunoResearch Laboratories (#115-545-003 and #111-605-003) and from Thermo Fisher Scientific (#A32728, #A32731 and #A31573). DAPI was obtained from Roth (#6335.1).

### Cell lines and cell culture

All urothelial carcinoma cell lines (UCCs) used in this research were kindly provided by Margaretha Skowron, Department of Urology, Heinrich Heine University Düsseldorf, Düsseldorf, Germany. K562 cells were obtained from the German Collection of Microorganisms and Cell Cultures GmbH (DMSZ, #ACC 10). Plat-E cells were kindly provided by Toshio Kitamura, Institute of Medical Science, University of Tokyo, Japan. Wild-type and *Atg13* KO MEFs were provided by Noboru Mizushima (University of Tokyo, Japan). Wild-type and *FIP200* KO U-2 OS cells were provided by Fulvio Reggiori (Aarhus University, Denmark). All cells were cultured in DMEM (Thermo Fisher Scientific/Gibco, #41965) or RPMI (Thermo Fisher Scientific/Gibco, #61870) supplemented with 10% FCS (Sigma-Aldrich #F0804, LOT BCCB7649 and #F9665, LOT 0001655439), 4.5 g/l D-glucose, 100 U/ml penicillin and 100 μg/ml streptomycin (Thermo Fisher Scientific/Gibco, #15140-122) at 37°C and 5% CO_2_ humidified atmosphere. During cell culture, cisplatin was added to the media of UCCs with every passage in concentrations of 2 µg/ml for 253J and 11 µg/ml for T24 cells. For experiments, cisplatin was removed unless specified.

### Cell viability assay

Cell viability was measured using either Alamar Blue or MTT assay. For both assays, 5,000-6,000 cells per well were seeded on a 96-well plate one day prior to the experiment. The following day, the cells were treated with different compounds as described above. For the Alamar Blue assay, 40 µM resazurin sodium salt (Cayman Chemicals, #14322) was added to the cells and incubated at 37°C for 1-4 h. Afterward, the absorbance was measured at 590 nm using a microplate reader (BioTek, Synergy Mx). For the MTT assay, 0.5 mg/ml MTT (Roth, #4022) was added to the cells and cells were incubated at 37°C for 1 h. Then the plates were centrifuged at 600 rcf for 5 min, and formazan crystals were dissolved in DMSO for 20 min in dark. The absorbance was measured at test (570 nm) and reference (650 nm) wavelengths using a microplate reader. The mean of the absorbance of untreated control samples was set as 100%.

### Cell death assay

Total cells treated with indicated stimuli were trypsinized and collected, then incubated in propidium iodide (PI) (5 μg/ml) solution at 4°C for 1 h. PI-positive cells were measured by flow cytometry (LSRFortessa, BD Biosciences).

### Caspase-3 activity assay

Cells treated with DMSO, VWK147 or STS were pelletized and quick-frozen in liquid nitrogen. Then, the cells were lysed with 50 µl of lysis buffer (20 mM HEPES, 84 mM KCl, 10 mM, MgCl_2_, 200 μM EDTA, 200 μM EGTA, 0.5% NP40, 1 μg/ml leupeptin, 1 μg/ml pepstatin, 5 μg/ml aprotinin) on ice for 10 min. The cell lysates then were transferred to a black microplate and mixed with 150 µl of reaction buffer (50 mM HEPES, 100 mM NaCl, 10% sucrose, 0.1% CHAPS, 2 mM CaCl_2_, 13.35 mM DTT, 70 µM Ac-DEVD-AMC). The fluorescence was measured for 2 h (Ex 360 nm, Em 450 nm) with every 2 min recording one data point using a microplate reader (CLARIOstar Plus, BMG LABTECH).

### Retroviral transduction

cDNA encoding human HSP70 was amplified from 253J cDNA by PCR using the following primers: fwd: GAAGGACCGAGCTCTTCTCG, rev: AGCAATCTTGGAAAGGCCCC. Subsequently, the HSP70 cDNA harboring 5′ and 3′ overlap to pMSCVhygro sequences was amplified and directly cloned into pMSCVhygro by sequence and ligation-independent cloning (SLIC) (63). The following primers were used: 3xFLAG-HSP70 fwd: GACGATGACAAGGGATCCGCCAAAGCCGCGGCGATCGGCA, 3xFLAG-HSP70 rev: CCCCTACCCGGTAGAATTCCTAATCTACCTCCTCAATGGTGGGGCC, pMSCVhygro fwd: GATTAGGAATTCTACCGGGTAGGGGAGGCGCTTTTCCCAAGGCA, and pMSCVhygro rev: CGGCTTTGGCGGATCCCTTGTCATCGTCATCCTTGTAATCGATG. Plat-E cells were used as packaging cells, and were transfected with the retroviral expression vectors pMSCVhygro, pMSCVhygro-3xFLAG-HSP70, pMSCVpuro or pMSCVpuro-mRFP-EGFP-rLC3 using FuGENE^®^ 6 (Roche, #11988387001). For the retroviral infection, pVSVG was co-transfected into Plat-E cells. After 48 h, UCCs were incubated with the corresponding retroviral supernatants containing 3 µg/ml Polybrene (Sigma-Aldrich, #H9268-106) for 72 h and subsequently selected in medium containing 200 µg/ml hygromycin or 1-2.5 µg/ml puromycin for about one week. After, T24 and T24-CR mRFP-EGFP-rLC3 expressing cells were additionally sorted for intermediate GFP signal at the Core Facility Flow Cytometry, University Clinic of Düsseldorf, in order to obtain a cell population expressing similar amounts of the protein.

### Immunoblotting

Cells were harvested by scraping and lysed in ice-cold lysis buffer (20 mM Tris/HCl, pH 7.5, 150 mM NaCl, 0.5 mM EDTA, 1% [v/v] Triton X-100, 1 mM Na_3_VO_4_, 10 μM Na_2_MoO_4_, 2.5 mM Na_4_P_2_O_7_, 10 mM NaF and protease inhibitor cocktail [Sigma-Aldrich, #P2714]). Alternatively, Na_3_VO_4_, Na_2_MoO_4_, Na_4_P_2_O_7_ and NaF were replaced by PHOSSTOP (Roche, #04906837001). Equal amounts of proteins were subjected to 8–15% SDS-PAGE. Proteins were then transferred to PVDF membranes (Millipore, #IPFL00010) or nitrocellulose membranes (Thermo Fisher Scientific, #88018) and analyzed using the indicated primary antibodies and appropriate IRDye-conjugated secondary antibodies. Protein signals were detected using an Odyssey Infrared Imaging system (LI-COR Biosciences) and quantified using Image Studio lite 4.0 and 5.2 (LI-COR Biosciences).

### Immunofluorescence

6.5-7.5 x 10^4^ cells were plated on glass coverslips one or two days prior to treatment. After treatment, the cells were fixed at room temperature using 4% paraformaldehyde for 15 min. Subsequently, the cells stably expressing mRFP-EGFP-rLC3 were directly stained with DAPI. The other cells were permeabilized with 50 µg/ml digitonin (Roth, #4005) for 5 min, blocked with 3% BSA (Roth, #8076)-PBS for 30 min, and then incubated with indicated primary antibodies for 2 h and corresponding secondary antibodies for 30 min. Cells were stained with DAPI and then embedded in ProLong Glass Antifade Mountant (Thermo Fisher Scientific, #P36980). The Zeiss Axio Observer 7 fluorescence microscope (Zeiss, Köln, Germany) with a Plan Apochromat 40x/1.4 oil objective (Zeiss, Köln, Germany) was used for imaging.

### Molecular dynamics simulations

To generate a conformational ensemble of the solvent-exposed hHSP90 protomer, a monomer taken from the cryo-EM structure of hHSP90 (PDB-ID 7L7I; (64)) was subjected to all-atom explicit solvent MD simulations. For this, the structure was prepared at a pH of 7.4 in Maestro (65) using the protein preparation wizard and PROPKA (66). MD simulations were performed with the Amber22 package of molecular simulation codes (67). The ff19SB force field (68) was used to parameterize the protein, Li-Merz parameters for ions (69), and the OPC water model (70) for the solvent. To generate input structures, a shell of at least 12 Å of OPC water, counter ions, and enough KCl to reach a salt concentration of 150 µM was added using Packmol-Memgen (71). To cope with long-range interactions, the Particle Mesh Ewald method (72) was used; the SHAKE algorithm (73) was applied to bonds involving hydrogen atoms, and a direct-space, non-bonded cut-off of 8 Å was used.

In the beginning, 17,500 steps of steepest descent and conjugate gradient minimization were performed; during 2500, 10,000, and 5000 steps, positional harmonic restraints with force constants of 25 kcal mol^-1^ Å^-2^, 5 kcal mol^-1^ Å^-2^, and zero, respectively, were applied to the solute atoms. Thereafter, 50 ps of NVT (constant number of particles, volume, and temperature) MD simulations were conducted to heat the system to 100 K, followed by 300 ps of NPT (constant number of particles, pressure, and temperature) MD simulations to adjust the density of the simulation box to a pressure of 1 atm and to heat the system to 300 K. During these steps, a harmonic potential with a force constant of 10 kcal mol^-1^ Å^-2^ was applied to the solute atoms. As the final step in thermalization, 300 ps of NVT-MD simulations were performed, while gradually reducing the restraint forces on the solute atoms to zero within the first 100 ps of this step. Afterward, twelve independent production runs of NVT-MD simulations with 1 µs length each were performed. For this, the starting temperatures of the MD simulations at the beginning of the thermalization were varied by a fraction of a Kelvin. Subsequently, the resulting runs were pooled and structures were extracted every ns. Then, a *k*-means clustering algorithm based on RMSD, as implemented in cpptraj (74), was used to assign the structures to five clusters, and the cluster representative was identified. After checking that all twelve runs were represented in the largest cluster, the representative of this cluster was chosen for subsequent docking experiments.

### Molecular docking

For the molecular docking, VWK147 was drawn and converted into a 3D structure with Maestro. Additional to the cluster representative from the above MD simulations, the X-ray crystal structures of hHSP90 with PDB ID 3Q6M (75) and 7L7I were used in their monomeric form and prepared with Maestro as described above. The ligand was subsequently docked into the binding pocket of the respective proteins using the combination of AutoDock as a docking engine and the DrugScore2018 distance-dependent pair-potentials as an objective function (76–78).

### Statistics

For immunoblotting, the density of each protein band was divided by the average of the density of all bands of this protein. The ratios were normalized to the loading control, and fold changes were calculated by dividing each normalized ratio by the average of the ratios of the solvent control (n ≥ 3). For PARP and Caspase-3, normalized bands representing the fragment of the protein of interest were additionally normalized to the normalized bands of the full-length protein. The results are shown as mean + standard deviation (SD) in the bar diagrams and in the dose-response curves. P values were either determined by ordinary one-way ANOVA with Dunnett’s post hoc or multiple comparisons test, by ordinary two-way ANOVA with Tukey’s multiple comparisons test or by unpaired t-test.

For cell viability assay, results are shown as mean ± standard deviation in the bar diagrams, and P values were determined by two-way ANOVA with Sidak’s post hoc test. All IC_50_ values were calculated using GraphPad Prism (versions 7.01 and 8.0.2). Compusyn 1.0 (34) was used to perform isobologram analysis for the combination of two compounds. The resulting combination index (CI) values represent the combination effects of two compounds and are interpreted as synergistic (CI ≤ 1), additive (CI = 1) or antagonistic (CI > 1) (34).

## Abbreviations

17-AAG: 17-Demethoxy-17-(2-propenylamino)geldanamycin
ATG: autophagy-related
BC: bladder cancer
CASM: conjugation of ATG8 proteins to single membranes
CI: combination index
EGFP: enhanced green fluorescent protein
HSF1: heat shock factor 1
HSP90: heat shock protein 90
HSR: heat shock response
KO: knockout
(MAP1)LC3: (microtubule-associated proteins 1A/1B) light chain 3
mRFP: monomeric red fluorescent protein
mTOR: mammalian target of rapamycin
IC_50_: half maximal inhibitory concentration
MTT: 3-(4,5-Dimethylthiazolyl-2)-2,5-diphenyl-2*H-*tetrazoliumbromid
PIK3C3: phosphatidylinositol 3-kinase catalytic subunit type 3
SQSTM1/p62: sequestosome 1
UCC: urothelial cell carcinoma
ULK1: unc-51-like autophagy activating kinase 1
VPS34: vacuolar protein sorting 34
WIPI: WD-repeat protein interacting with phosphoinositides
WT: wild-type

