## supporting information for "Small-molecule inhibitor of C-terminal HSP90 dimerization modulates autophagy and functions synergistically with mTOR inhibition to kill cisplatin-resistant cancer cells"

#shared first authorship

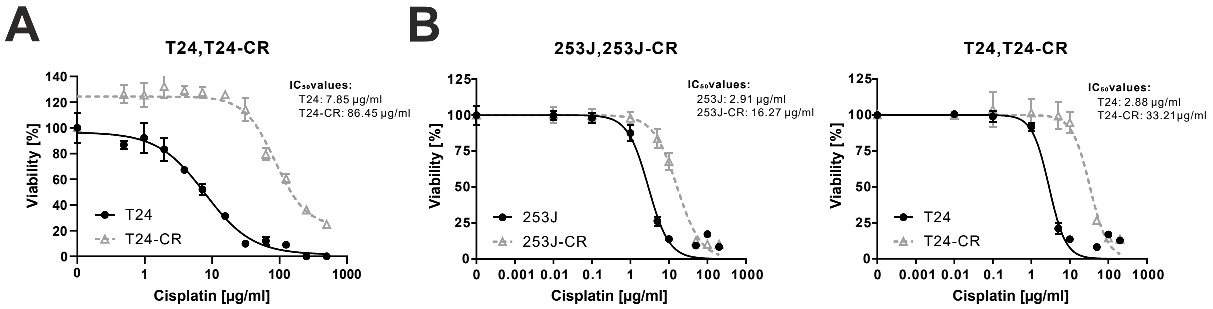


**Suppl. Figure S1: Characterization of cisplatin-sensitive and -resistant urothelial bladder carcinoma cell lines.** 253J, 253J-CR, T24 and T24-CR urothelial carcinoma cells were treated with indicated concentrations of cisplatin for (**A**) 24 or (**B**) 72 h. After treatment, cell viability was measured using Alamar Blue assay. Results are shown as means ± SD of (**A**) one to (**B**) three independent experiments performed in triplicates for each treatment.

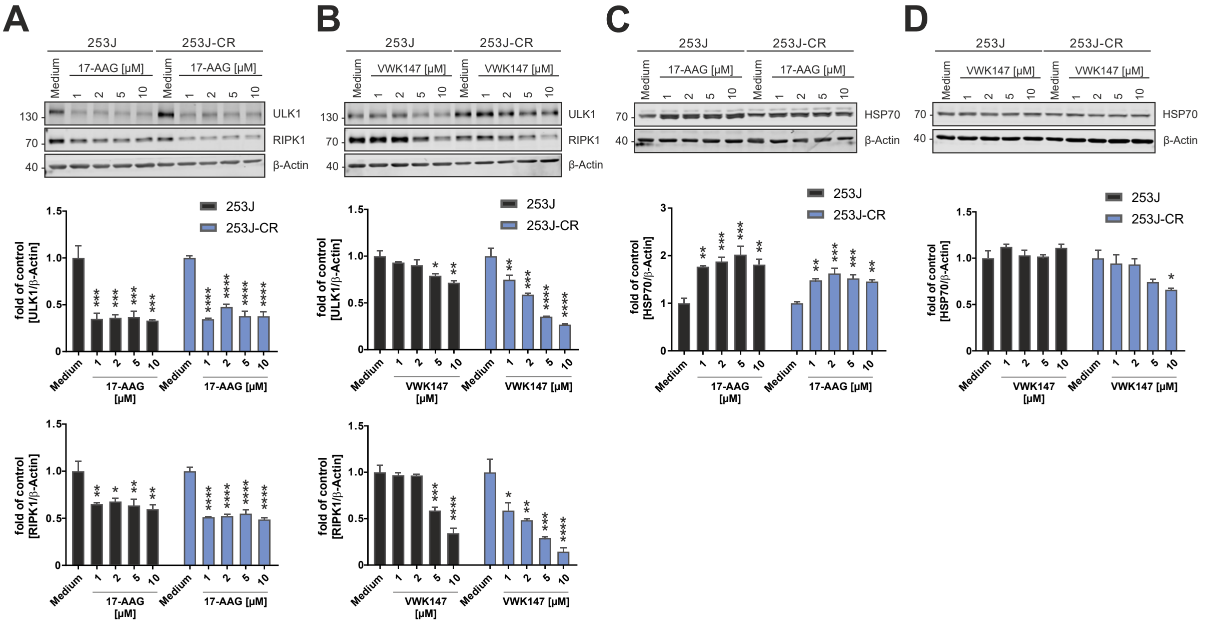


**Suppl. Figure S2: VWK147 destabilizes HSP90 clients but does not induce a heat-shock response.** (**A-D**) 253J and 253J-CR urothelial carcinoma cells were treated with the indicated concentrations of 17-AAG or VWK147 for 6 h. After treatment, the cells were lysed, and cellular lysates were immunoblotted for ULK1, RIPK1, Actin, and HSP70, respectively. One representative immunoblot is shown. The quantifications of indicated ratios are from three independent experiments (means + SD). P values were determined by ordinary one-way ANOVA with Dunnett´s post hoc test (comparison to the solvent control of the respective cell line). *p ≤ 0.05; **p ≤ 0.01; ***p ≤ 0.001; ****p ≤ 0.0001.

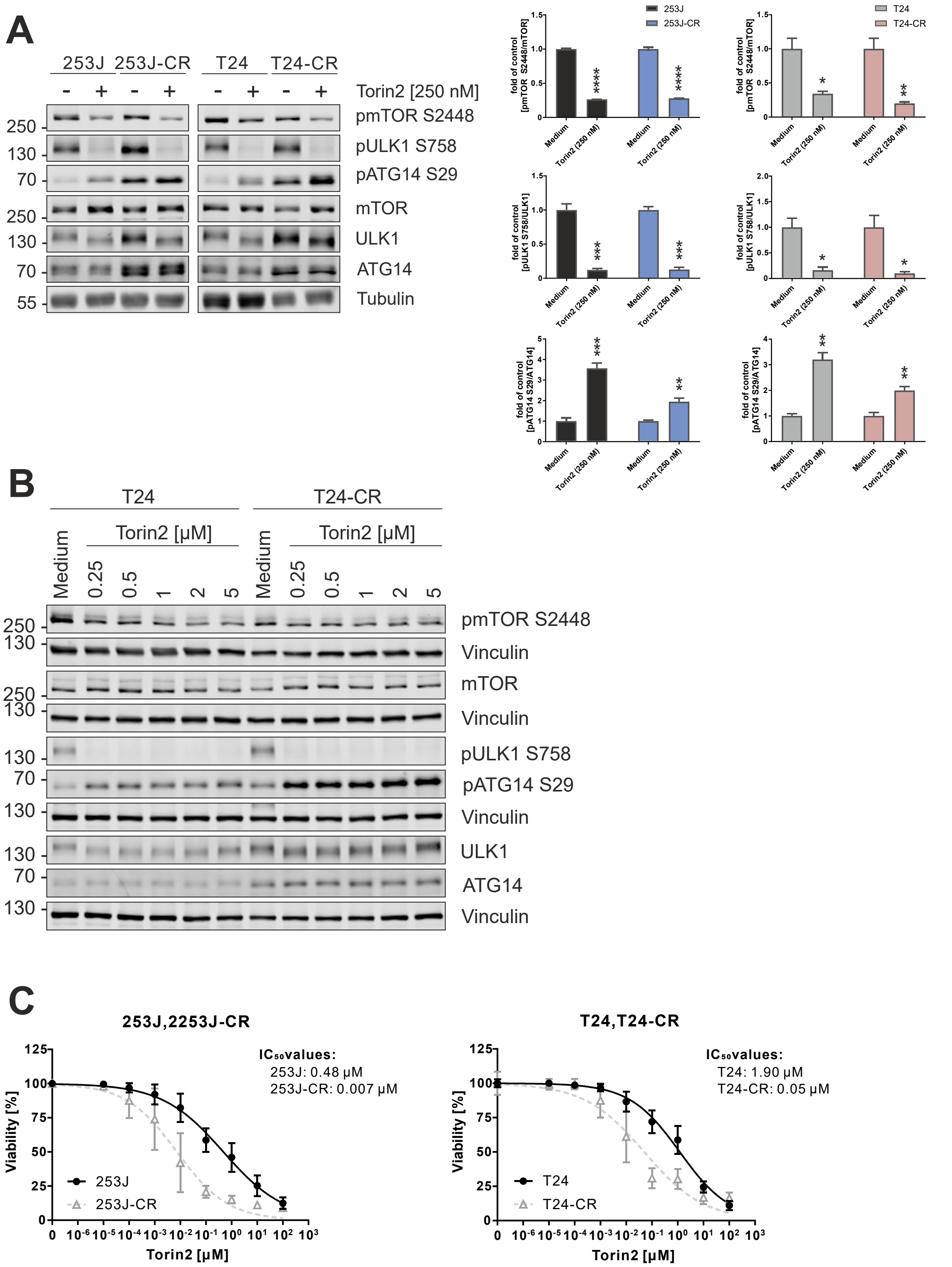


**Suppl. Figure S3: Characterization of Torin2 efficacy in urothelial bladder carcinoma cell lines.** (**A**) 253J, 253J-CR, T24 and T24-CR urothelial carcinoma cells were treated with 250 nM Torin2 for 2 h. After treatment, the cells were lysed, and cellular lysates were immunoblotted for phospho-mTOR

Ser2448, mTOR, phospho-ULK1 Ser758, ULK1, phospho-ATG14 Ser29, ATG14 and tubulin. One representative immunoblot is shown. The quantifications of indicated ratios are from three independent experiments (means + SD). P values were determined by unpaired t test (comparison to the solvent control of the respective cell line). $^{*}p \leq 0.05$; $^{**}p \leq 0.01$; $^{***}p \leq 0.001$; $^{****}p \leq 0.0001$. (**B**) T24 and T24-CR urothelial carcinoma cells were treated with indicated concentrations of Torin2 for 24 h. After treatment, the cells were lysed, and cellular lysates were immunoblotted for phospho-mTOR Ser2448, mTOR, phospho-ULK1 Ser758, ULK1, phospho-ATG14 Ser29, ATG14 and Vinculin (n=1). (**C**) 253J, 253J-CR, T24 and T24-CR urothelial carcinoma cells were treated with indicated concentrations of Torin2 for 72 h. After treatment, cell viability was measured using Alamar Blue assay. Results are shown as means ± SD of three independent experiments performed in triplicates for each treatment.

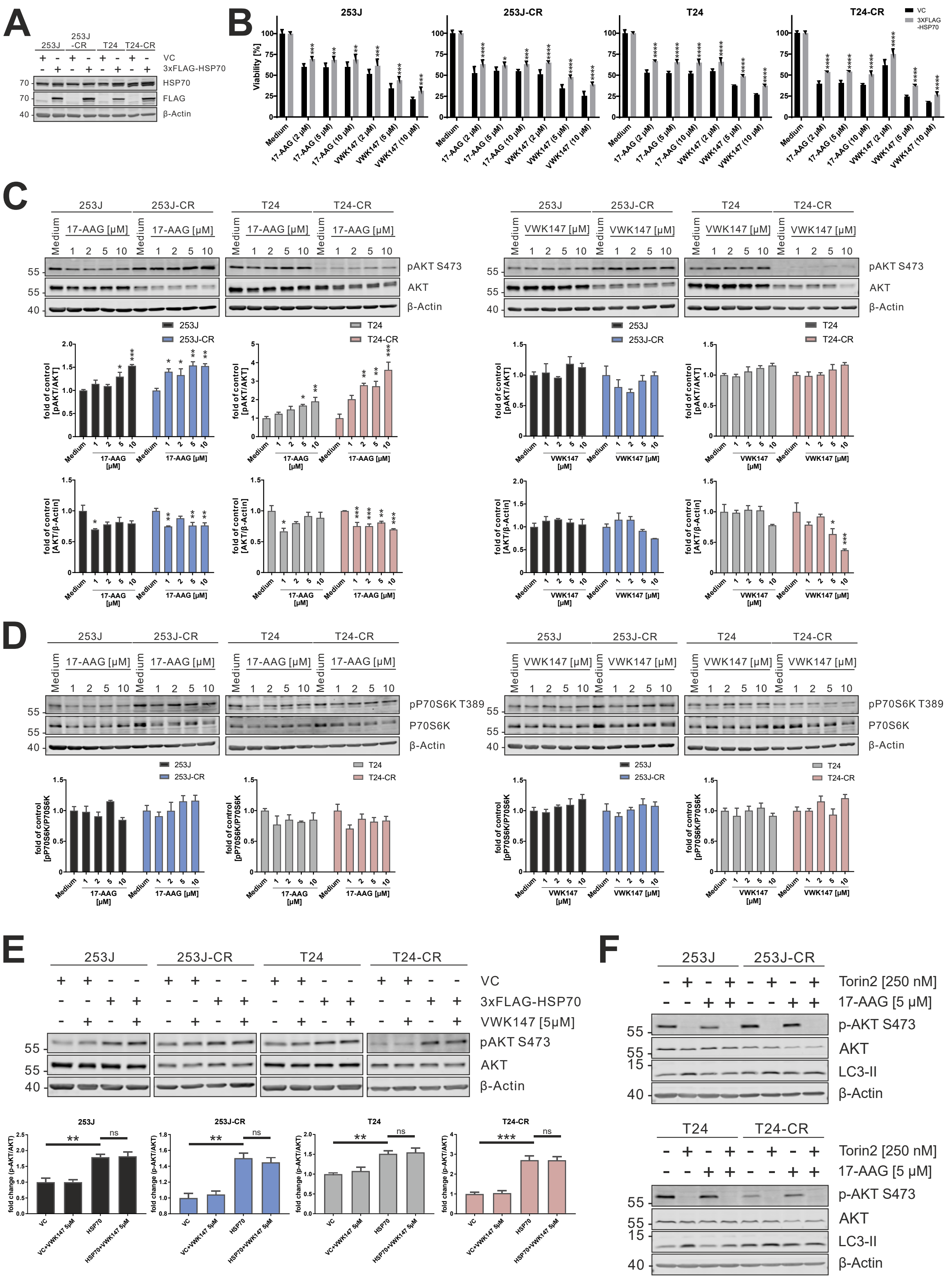


**Suppl. Figure S4: Forced expression of HSP70 reduces cytotoxic effect of VWK147.** (**A**) 253J, 253J-CR, T24 and T24-CR urothelial carcinoma cells transfected with either empty vector (VC) or cDNA encoding 3xFLAG-HSP70 were lysed and cellular lysates were immunoblotted for HSP70, FLAG and β-Actin. One representative immunoblot is shown. (**B**) 253J, 253J-CR, T24 and T24-CR urothelial carcinoma cells transfected with either empty vector (VC) or cDNA encoding 3xFLAG-HSP70-were treated with indicated concentrations of 17-AAG or VWK147 for 24 h. After treatment, cell viability was measured using MTT assay. Results are shown as means + SD of three independent experiments performed in triplicates for each treatment. P values were determined by ordinary two-way ANOVA with Sidak´s post hoc test. *p ≤ 0.05; **p ≤ 0.01; ***p ≤ 0.001; ****p ≤ 0.0001 (**C**) 253J, 253J-CR, T24 and T24-CR cells-were treated with indicated concentrations of 17-AAG or VWK147 for 6 h. After treatment, the cells were lysed, and cellular lysates were immunoblotted for phospho-AKT Ser473, AKT and β-Actin. One representative immunoblot is shown. The quantifications of indicated ratios are from three independent experiments (means + SD). P values were determined by ordinary one-way ANOVA with Dunnett´s post hoc test (comparison to the solvent control of the respective cell line). *p ≤ 0.05; **p ≤ 0.01; ***p ≤ 0.001; ****p ≤ 0.0001 (**D**) 253J, 253J-CR, T24 and T24-CR cells were treated with indicated concentrations of 17-AAG or VWK147 for 6 h. After treatment, the cells were lysed, and cellular lysates were immunoblotted for phospho-p70/S6K Thr389, p70/S6K and β-Actin. One representative immunoblot is shown. The quantifications of indicated ratios are from three independent experiments (means + SD). Significance was analyzed by ordinary one-way ANOVA with Dunnett´s post hoc test (comparison to the solvent control of the respective cell line). (**E**) 253J, 253J-CR, T24 and T24-CR cells transfected with either empty vector (VC) or cDNA encoding 3xFLAG-HSP70-were treated with 5 µM VWK147 for 24 h. After treatment, the cells were lysed, and cellular lysates were immunoblotted for phospho-AKT Ser473, AKT and β-Actin. One representative immunoblot is shown. The quantifications of indicated ratios are from three independent experiments (means + SD). P values were determined by ordinary one-way ANOVA with Tukey´s post hoc test. **p ≤ 0.01; ***p ≤ 0.001; NS, non-significant. (**F**) 253J, 253J-CR, T24 and T24-CR urothelial carcinoma cells were treated with indicated mono- or combination treatments (250 nM Torin2, 5 µM 17-AAG) for 6 h. After treatment, the cells were lysed, and cellular lysates were immunoblotted for phospho-AKT Ser473, AKT, LC3 and β-Actin. One representative immunoblot is shown.

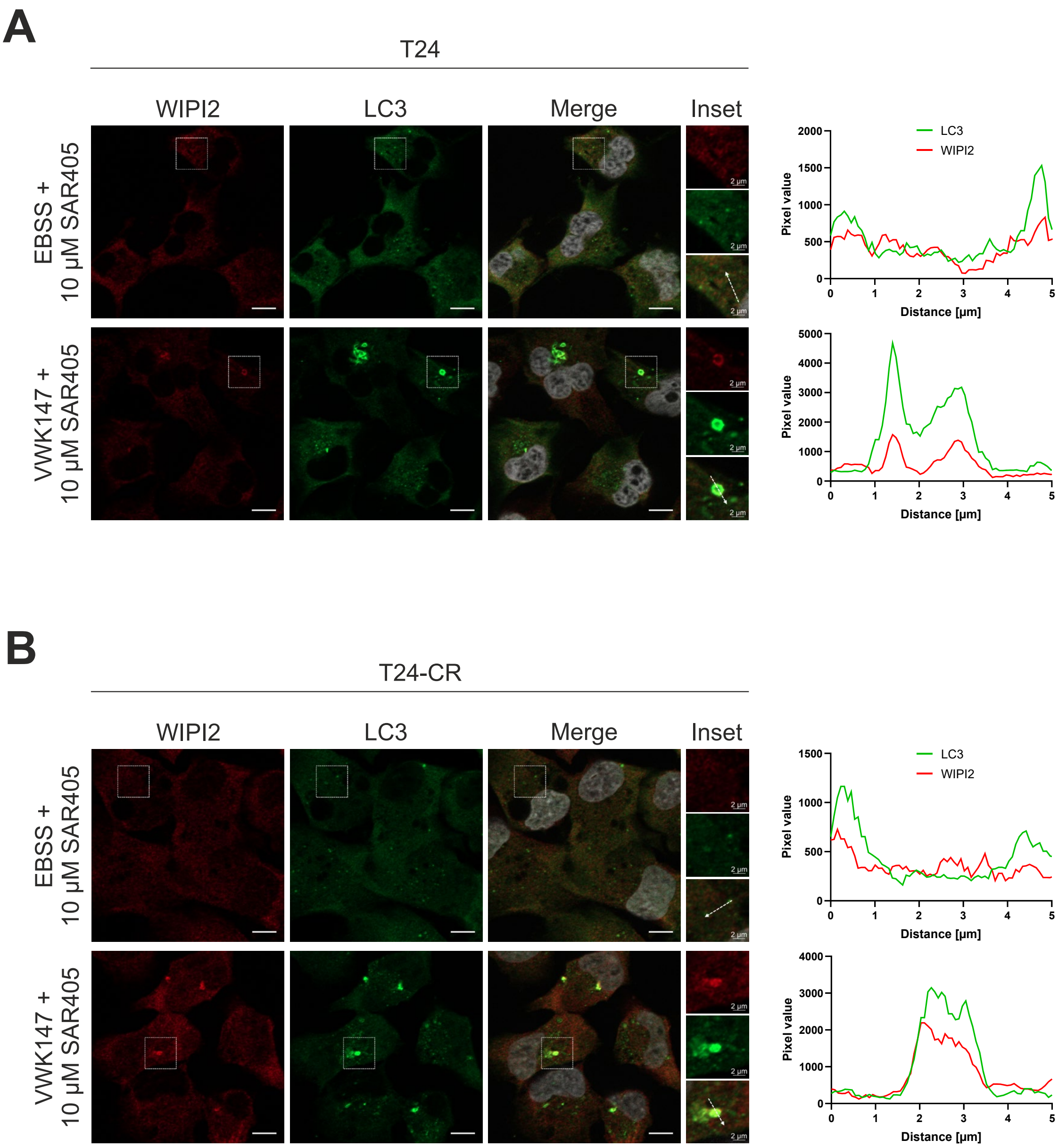


**Suppl. Figure 5: VWK147 induces LC3-WIPI2 aggregates in T24 and T24-CR urothelial carcinoma cells.** (**A**) T24 and (**B**) T24-CR urothelial carcinoma cells were grown on glass coverslips one or two days prior to treatment. Cells were treated with 10 µM SAR405 in combination with EBSS or 5 µM VWK147 for 4 h. Imaging was performed using a Zeiss Axio Observer 7 fluorescence microscope equipped with a 40x/1.4 Oil DIC M27 Plan-Apochromat objective and ApoTome 2. Representative sections are depicted. Scale bars: 10 µM and 2 µM. The line graphs represent the pixel intensities of the areas indicated by the respective dashed white arrows shown in the insets.

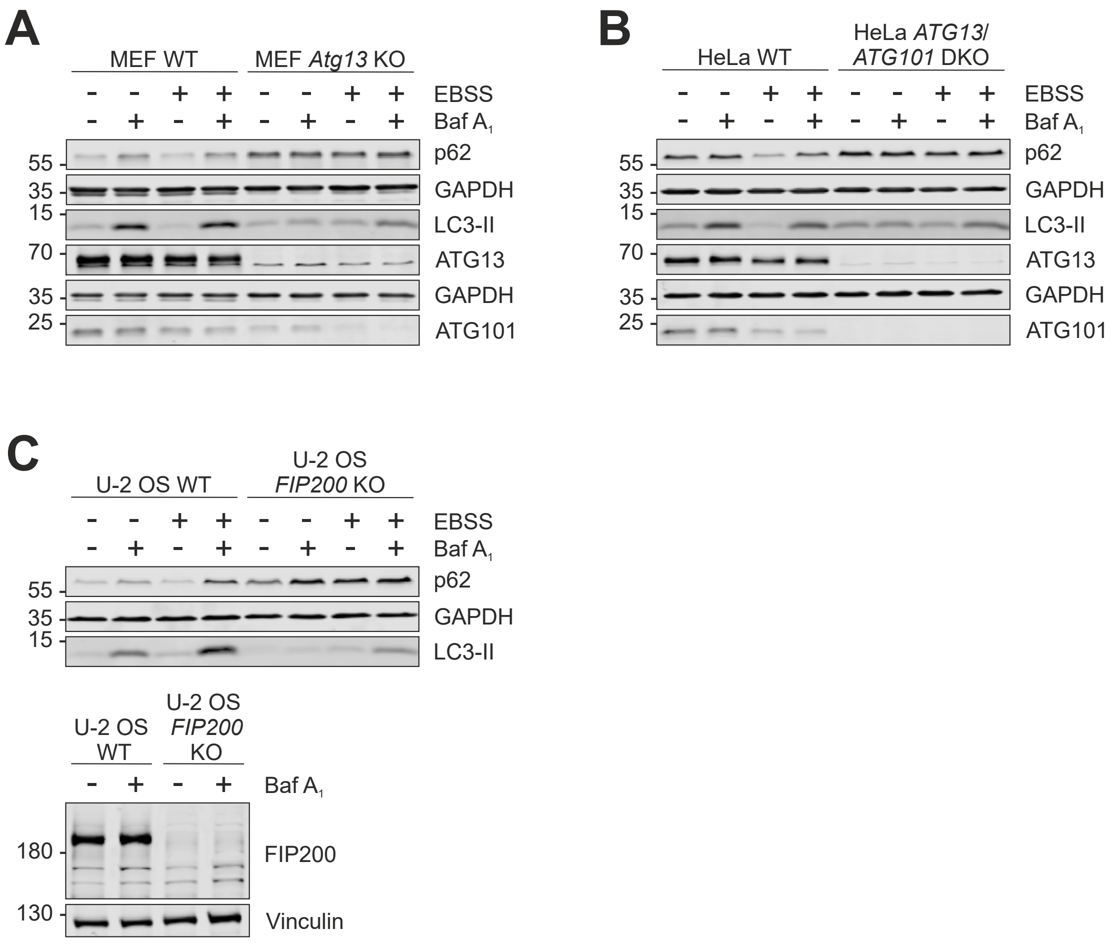


**Suppl. Figure 6: Knockout of *Atg13/ATG13*, *ATG101* or *FIP200* severely impairs canonical autophagy in MEF, HeLa and U-2 OS cells.** (**A**) MEF WT and MEF *Atg13* KO, (**B**) HeLa WT and HeLa *ATG13*/*ATG101* DKO and (**C**) U-2 OS WT and U-2 OS *FIP200* KO cells were treated with mono- or combination treatments (EBSS, 20 nM Bafilomycin $A_1$) for 6 h. After treatment, the cells were lysed, and cellular lysates were immunoblotted for p62, LC3, GAPDH, FIP200, ATG13, ATG101 and Vinculin (n=1).

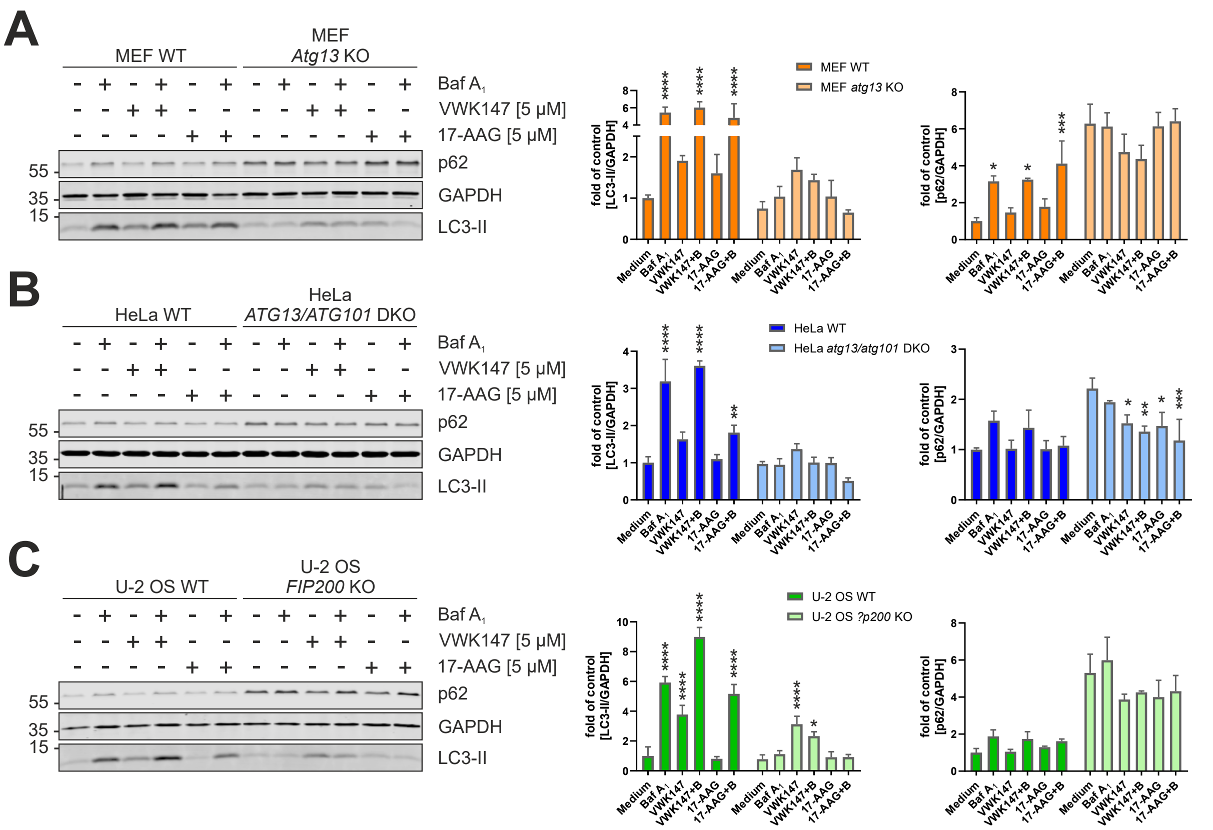


**Suppl. Figure 7: 17-AAG does not induce non-canonical autophagy.** (**A**) MEF WT and MEF *Atg13* KO, (**B**) HeLa WT and HeLa *ATG13/ATG101* DKO and (**C**) U-2 OS and U-2 OS *FIP200* KO were treated with 5 µM VWK147 or 17-AAG in presence or absence of 20 nM bafilomycin $A_1$ for 6 h. After treatment, the cells were lysed, and cellular lysates were immunoblotted for p62, LC3 and GAPDH. One representative immunoblot is shown. The quantifications of indicated ratios are from three independent experiments (means + SD). P values were determined by ordinary two-way ANOVA with Tukey´s multiple comparisons test (comparison to the solvent control of the respective cell line). *p ≤ 0.05; **p ≤ 0.01; ***p ≤ 0.001; ****p ≤ 0.0001.

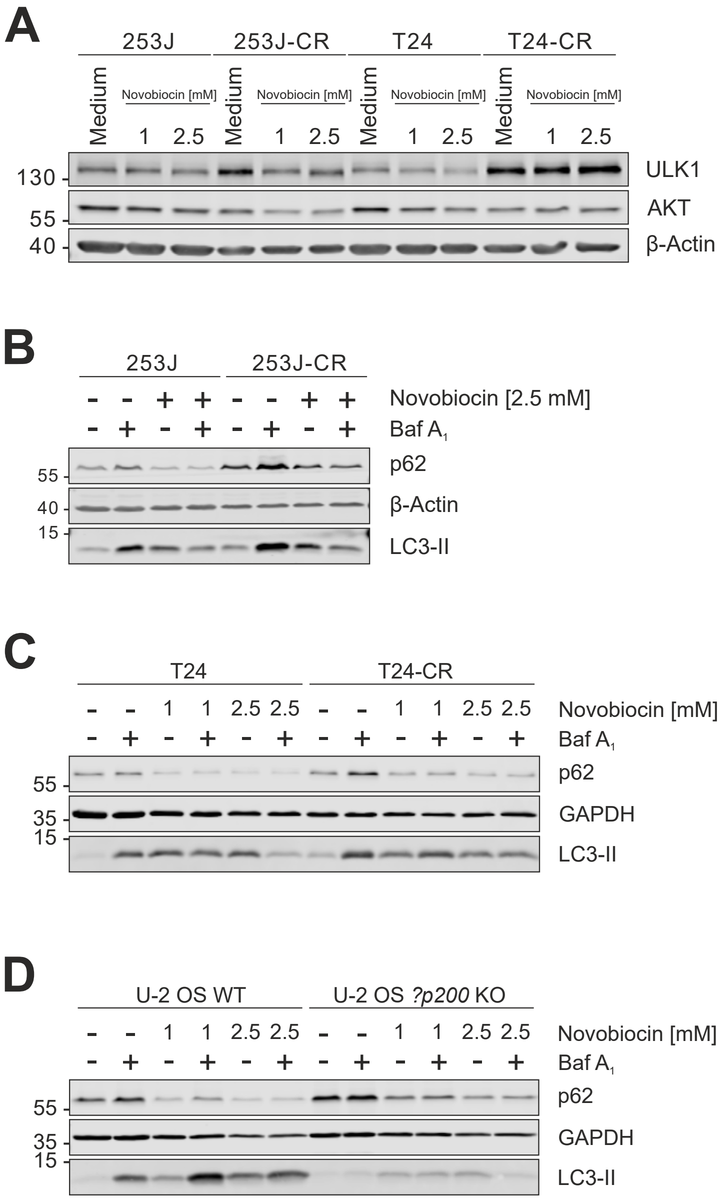


**Suppl. Figure 8: Novobiocin induces non-canonical LC3 lipidation.** (**A**) 253J, 253J-CR, T24 and T24-CR urothelial carcinoma cells were treated with indicated concentrations of Novobiocin for 6 h. After treatment, the cells were lysed, and cellular lysates were immunoblotted for ULK1, AKT and β-Actin. One representative immunoblot is shown. (**B**) 253J, 253J-CR, (**C**) T24 and T24-CR urothelial carcinoma cells as well as (**D**) U-2 OS WT and U-2 OS *FIP200* KO osteosarcoma cells were treated with indicated mono- or combination treatments (1 mM or 2.5 mM Novobiocin, 20 nM bafilomycin $A_1$) for 6 h. After treatment, the cells were lysed, and cellular lysates were immunoblotted for p62, LC3 and β-Actin. One representative immunoblot is shown (n=3).
